# Potent neutralization of Marburg virus by a vaccine-elicited monoclonal antibody

**DOI:** 10.1101/2025.05.14.654121

**Authors:** Amin Addetia, Lisa Perruzza, Young-Jun Park, Matthew McCallum, Cameron Stewart, Jack T. Brown, Alessia Donati, Katja Culap, Alessio Balmelli, Michal Gazi, Ricardo Carrion, Davide Corti, Fabio Benigni, David Veesler

## Abstract

Marburg virus (MARV) is a filovirus that causes a severe and often lethal hemorrhagic fever. Despite the increasing frequency of MARV outbreaks, no vaccines or therapeutics are licensed for use in humans. Here, we designed mutations that improve the expression and thermostability of the prefusion MARV glycoprotein (GP) ectodomain trimer, which is the sole target of neutralizing antibodies and vaccines in development. We discovered a fully human monoclonal antibody, MARV16, that broadly neutralizes all MARV isolates as well as Ravn virus and Dehong virus with 40 to 100-fold increased potency relative to previously described antibodies. We determined a cryo-electron microscopy structure of MARV16-bound MARV GP showing that MARV16 recognizes a prefusion-specific epitope spanning GP1 and GP2, blocking receptor binding and preventing conformational changes required for viral entry. We further reveal the architecture of the MARV GP glycan cap, which shields the receptor binding site (RBS), underscoring architectural similarities with distantly related filovirus GPs. MARV16 and previously identified RBS-directed antibodies can bind MARV GP simultaneously, paving the way for a MARV therapeutic antibody cocktail. MARV GP stabilization along with the discovery of a potent neutralizing antibody will advance treatment and prevention options for MARV.

## Introduction

Marburg virus (MARV) belongs to the *Filoviridae* family and causes Marburg virus disease (MVD), which is characterized by a hemorrhagic fever with a case fatality rate ranging from 24 to 88% (*1–3*). In recent years, multiple African countries have experienced MARV outbreaks including Ghana in 2022, Equatorial Guinea and Tanzania in 2023, and Rwanda in 2024 (*4–7*). Another outbreak is currently ongoing in Tanzania. Although recent outbreaks were contained, larger MARV outbreaks are likely to occur, similar to the 2013-2016 outbreak of the related Ebola virus (EBOV) in West Africa which led to greater than 28,600 infections and 11,325 casualties (*8*, *9*). The recurring and frequent spillovers of MARV underscores the necessity for licensed vaccines and therapeutics, which are currently not available.

Vaccines and monoclonal antibodies in development against MARV target the viral glycoprotein (GP) as GP-directed antibodies have been suggested to be the primary correlate of protection against MVD (*10*). MARV GP is a homotrimeric protein anchored in the viral membrane and is responsible for recognition of the host receptor, NPC1, and membrane fusion, leading to viral entry (*11–13*). The GP is proteolytically cleaved by furin during viral morphogenesis to produce the GP1 and GP2 subunits that remain covalently linked by a disulfide bond (*14–16*). GP1 contains three domains: core, glycan cap, and mucin-like domain. The GP1 core contains the receptor-binding site (RBS), which is shielded from neutralizing antibodies by the highly glycosylated glycan cap and mucin-like domain (*11*, *15*, *17*, *18*). Unlike for the EBOV or Sudan virus (SUDV) GPs, an ordered glycan cap has not been visualized for MARV GP, suggesting that the RBS might be more exposed for MARV GP compared to the EBOV and SUDV GPs (*15*, *16*, *18*, *19*). GP2 is also composed of three domains: wing, core, and transmembrane domain. The GP2 core is the fusion machinery that promotes fusion of the viral and host membranes (*20–22*). The wing domain is unique to the *Marburgvirus* genus and wraps around the GP equator, which was proposed to limit recognition by neutralizing antibodies (*16*).

Several MARV investigational vaccines have advanced to Phase I or II clinical trials with one of them, cAd3-Marburg, being deployed during the MARV outbreak in Rwanda in 2024 (*7*, *10, 23–27*). These vaccines either display MARV GP on a viral vector or encode for MARV GP. These vaccines may benefit from the identification and inclusion of mutations that increase GP expression and prefusion stability, similar to the stabilizing mutations that were incorporated into the SARS-CoV-2 and RSV vaccines (*28–32*). Only one stabilizing mutation that promotes trimer formation of the MARV GP ectodomain has been identified to date (*33*). In contrast, several stabilizing mutations have been identified for the EBOV and SUDV GP, suggesting that further optimization of prefusion MARV GP may be possible (*33*, *34*).

Monoclonal antibodies targeting multiple MARV GP domains have been isolated from MARV survivors or GP-immunized animals (*35–38*). However, only antibodies targeting the RBS had detectable, albeit weak, neutralizing activity against MARV (*35*). Several of these RBS-directed antibodies show protective efficacy in animal models with one of them, MR191, being the precursor to the investigational therapeutic monoclonal antibody MBP01 (*35*, *39*, *40*). Given that the neutralization potency of RBS-directed antibodies, including MR191, can be reduced by single mutations within the highly variable glycan cap (*35*), an antibody cocktail may prove to be more resilient to viral evolution.

Here, we set out to identify MARV GP stabilizing mutations to develop an immunogen allowing for subsequent discovery of vaccine-elicited antibodies that potently neutralize MARV. We identified two mutations in the MARV GP2 heptad repeat 1-C (HR1_C_) within the GP2 core domain that increase both the expression and thermostability of the prefusion ectodomain trimer while retaining its native prefusion structure and antigenicity. Immunization of a humanized transgenic mouse with prefusion-stabilized MARV GP ectodomain trimer enabled isolation of a potent neutralizing antibody, designated MARV16, that targets a prefusion GP quaternary epitope spanning the GP1 and GP2 subunits. We further demonstrate that MARV16 potently neutralizes historical and contemporary MARV variants as well as the related Ravn (RAVV) and Dehong viruses (DEHV). Our structure resolves a portion of the previously elusive MARV GP1 glycan cap, indicating that it partially shields the RBS, as is the case for EBOV GPs. Finally, we show that MARV16 can bind to MARV GP simultaneously with RBS-directed antibodies, providing a pathway toward developing a therapeutic antibody cocktail for MVD.

## Results

### Expression and characterization of the MARV GPΔMuc ectodomain

In order to produce a soluble, prefusion MARV GP ectodomain trimer, we first designed a MARV GP construct lacking the mucin-like (residues 257-425) and transmembrane domains (residues 638-681) **(Figure 1A)**. The mucin-like domain was omitted from our construct as antibodies targeting this domain are unlikely to be neutralizing (*41*, *42*). We additionally incorporated three previously described mutations, W439A, F445G, and F447N, that increase cleavage of precursor GP by furin (*15*) along with the H589I mutation that promotes trimer formation in the absence of exogenous trimerization domain (*33*). Co-expression of this construct, termed MARV GPΔMuc, with furin yielded monodisperse trimers **(Figure 1B)** as visualized by negative stain electron microscopy. Binding to MR191 **(Figure 1C)** confirmed proper folding and antigenicity of our MARV GPΔMuc ectodomain, which we used for a subsequent antibody discovery campaign.

**Figure 1.**
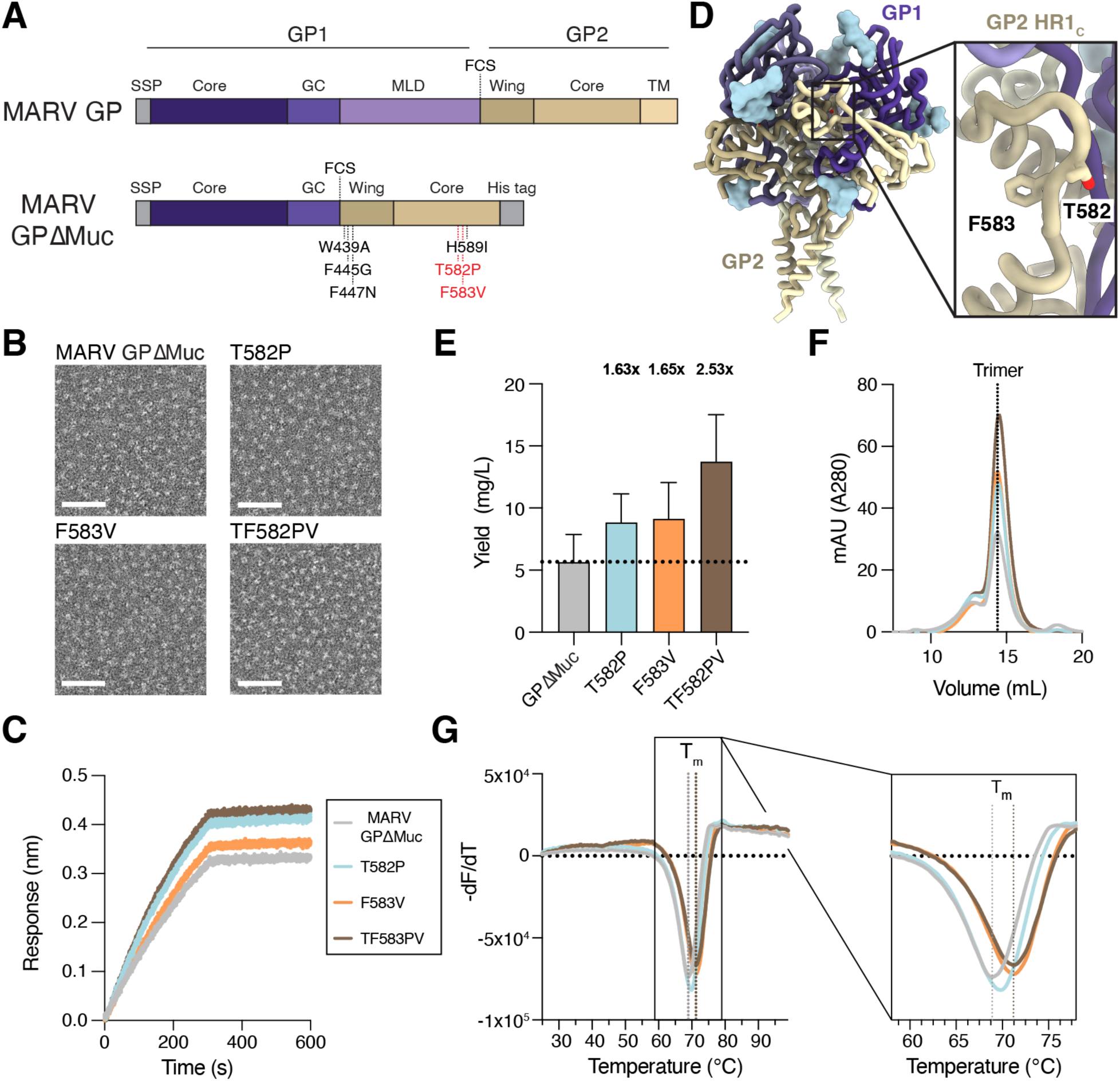
Computational design of MARV GP prefusion-stabilizing mutations. **A)** Schematic of the MARV GP domain organization and engineered mutations included in the MARV GPΔMuc ectodomain construct. GP2 HR1_C_ residue mutations introduced to further stabilize the MARV GPΔMuc ectodomain are shown in red. FCS: furin cleavage site. **B)** Representative negative stain electron microscopy micrograph of the original and stabilized MARV GPΔMuc ectodomains. Scale bar: 50 nm. **C)** Binding of the original and stabilized MARV GPΔMuc ectodomain to immobilized MR191 IgG as assessed by biolayer interferometry (BLI). Data presented are from one representative out of four biological replicates. **D)** Ribbon diagram of the Ravn virus GP structure (PDB: 6BP2) highlighting the residues mutated in GP2 HR1_c_. GP1 is shown in purple, GP2 is shown in gold, and N-linked glycans are rendered as light blue surfaces. **E)** Recombinant production yields of the original and stabilized MARV GPΔMuc ectodomains purified from transiently transfected Expi293 cells. The fold-increase in yield of the stabilized MARV GPΔMuc ectodomains relative to the original ectodomain is displayed above the plot. The bar represents the mean yield and the error bar indicates the standard deviation across four biological replicates. **F)** Size-exclusion chromatograms of the original and stabilized MARV GPΔMuc. Data presented are from one biological replicate and are representative of three additional biological replicates. **G)** Differential scanning fluorimetry analysis of the original and stabilized MARV GPΔMuc ectodomains. Data are shown as the first negative derivative of the fluorescence intensity with respect to temperature. The melting temperature (T_m_) of the original MARV GPΔMuc and MARV GPΔMuc TF582PV ectodomain are indicated with the dotted vertical lines. Data presented are from one biological replicate and are representative of three additional biological replicates. Six technical replicates were conducted and averaged for each biological replicate.

### Identification of stabilizing mutations in MARV GP2 HR1_C_

Prior studies on EBOV GP have identified mutations in GP2 HR1_C_ that improve expression yields with and without fusion of a trimerization domain (*33*, *34*). However, porting the EBOV GP T578P stabilizing mutation to MARV GP did not yield the same expression enhancement (*33*). We used ProteinMPNN (*43*) to assist identification of MARV GP HR1_C_ mutations that could improve expression of the prefusion ectodomain trimer **(Figure 1A and Figure 1D)**. We found that the T582P and F583V individual residue substitution respectively improved expression yields 1.6-fold and 1.7-fold, compared to our original MARV GPΔMuc, and led to a 2.5-fold enhancement when combined together **(Figure 1E)**. All three designed MARV GPΔMuc ectodomain constructs eluted as monodisperse species and at a similar retention volume to MARV GPΔMuc when analyzed by size-exclusion chromatography **(Figure 1F)**. Electron microscopy imaging of negatively stained samples confirmed the monodispersity of the constructs **(Figure 1B)** and retention of MR191 binding **(Figure 1C)** indicated that introduction of the HR1_c_ mutations did not alter the conformational integrity or antigenicity of MARV GP. Both the T582P (T_m_: 69.7 ± 0.2°C) and F583V (T_m_: 71.2 ± 0.1°C) mutations improved the thermostability of MARV GPΔMuc, increasing the melting temperature (T_m_) by 0.9 and 2.4°C, respectively. The TF582PV double (designated MARV GPΔMuc_PV_) mutant was endowed with a 2.6°C greater melting temperature than the original MARV GPΔMuc (T_m_: 71.4 ± 0.2 versus 68.8 ± 0.3°C), suggesting improved stability of the prefusion state **(Figure 1G)**.

### Discovery of a potent MARV neutralizing antibody

All monoclonal antibodies identified to date that target MARV GP display no or weak neutralization potency against MARV GP pseudoviruses and even weaker, if any, activity against authentic MARV (*35–38*). To identify potent MARV neutralizing antibodies, we immunized Alloy ATX transgenic mice harboring the human immunoglobulin loci encoding for the heavy chain and either the lambda (ATX-GL) or the kappa (ATX-GK) light chain. Two ATX-GL and two ATX-GK mice were immunized with prefusion-stabilized MARV GPΔMuc for a total of 3 doses **(Figure 2A)**. Mice were sacrificed 6 days after the last boost, and peripheral blood, spleen, and lymph nodes (LNs) were collected and cells freshly isolated. MARV GPΔMuc-specific memory B cells were selected via fluorescence-assisted cell sorting and variable domain (VH/VL) sequences subsequently retrieved by reverse-transcription PCR (RT-PCR).

**Figure 2.**
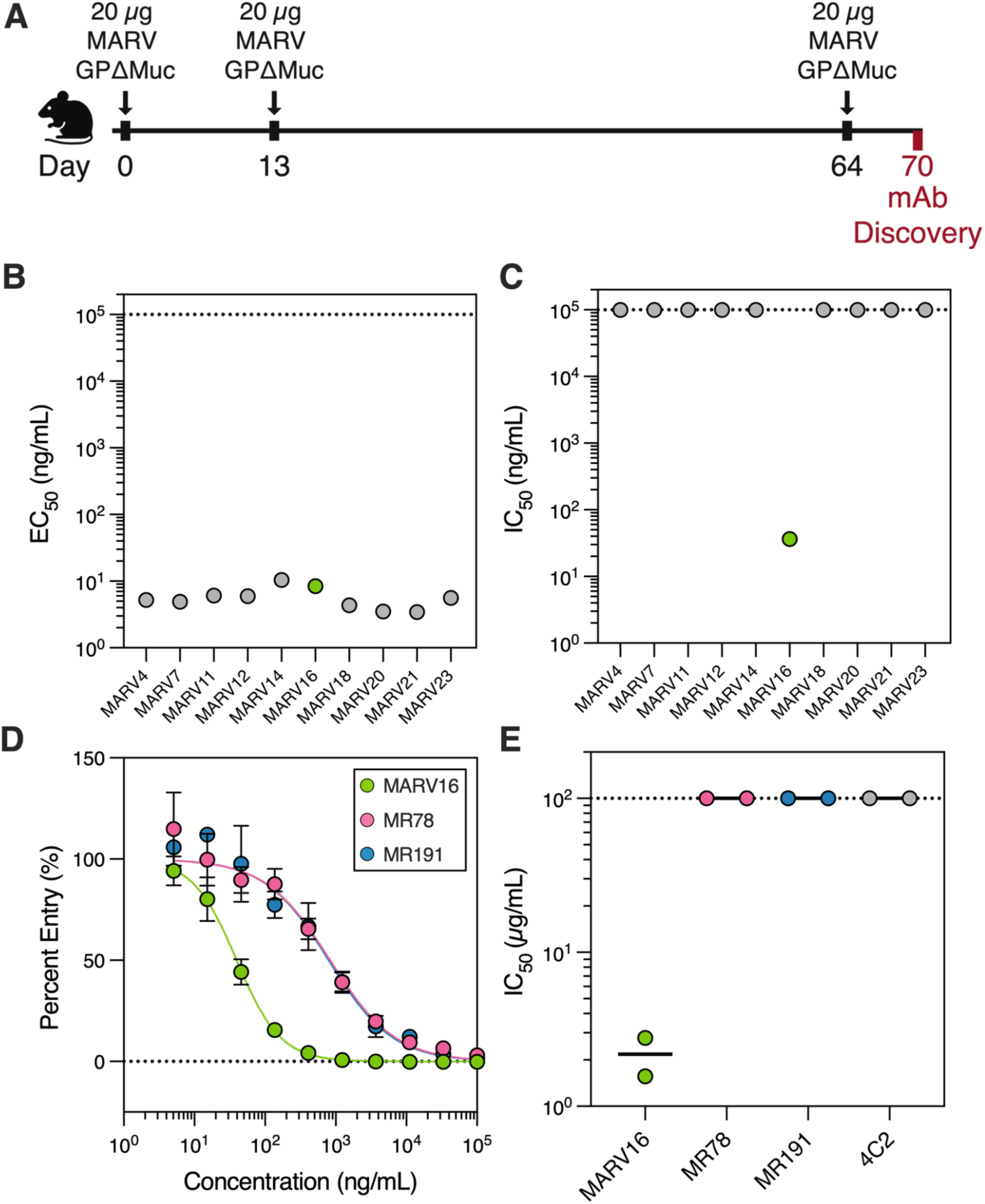
Discovery of a potent fully human MARV neutralizing antibody. **A)** Schematic of the immunization schedule used to discover MARV GP-directed monoclonal antibodies. **B)** Binding of the 10 monoclonal antibodies isolated in this study to the MARV GPΔMuc ectodomain as measured by ELISA. Data points represent averaged half-maximal effective concentrations (EC_50_) values obtained from two biological replicates conducted with distinct batches of protein. Each biological replicate was conducted in technical duplicate. **C)** Evaluation of neutralizing activity of the 10 monoclonal antibodies against VSV pseudotyped with the vaccine-matched MARV/Musoke GP. Data points represent averaged half-maximal inhibitory concentrations (IC_50_) values obtained from at least two biological replicates using distinct batches of pseudovirus and monoclonal antibodies. Each biological replicate was conducted with three technical replicates. **D)** Dose-response neutralization curves for MARV16, MR78, and MR191 against VSV pseudotyped with the MARV/Musoke GP. Data reflect a single biological replicate conducted with three technical replicates and are representative of three to five additional biological replicates. **E)** Neutralization potency of MARV16, MR78, MR191, and 4C2, a MERS-CoV spike protein-directed antibody, against authentic MARV/Musoke. Data points reflect 50% plaque reduction neutralization test [PRNT_50_] values obtained from two biological replicates using distinct batches of monoclonal antibodies. The black line indicates the mean PRNT_50_ value obtained from the two biological replicates.

We recovered 10 antibodies that bound to MARV GPΔMuc (half-maximal effective concentration [EC_50_]: 3.4 - 10.4 ng/mL) **(Figure 2B; Figure S1)** and one of them, designated MARV16, potently neutralized VSV pseudotyped with the vaccine-matched MARV/Musoke GP (half-maximal inhibitory concentration [IC_50_]: 36.4 ng/mL) **(Figure 2C; Figure S1)**. MARV16 is 42-fold and 39-fold more potent than the previously described RBS-directed neutralizing antibodies (*35*) MR78 (IC_50_: 1,520 ng/mL) and MR191 (IC_50_: 1,407 ng/mL), respectively, as measured side-by-side using MARV/Musoke GP VSV pseudovirus **(Figure 2D)**. Furthermore, MARV16 neutralized authentic MARV/Musoke (50% plaque reduction neutralization test [PRNT_50_]: 2.2 µg/mL) with >45-fold potency than MR78 (PRNT_50_: >100 µg/mL) and MR191 (PRNT_50_: >100 µg/mL) **(Figure 2E, Figure S2)**, respectively, establishing MARV16 as a best-in-class MARV neutralizing antibody.

We assessed the kinetics and affinity of the MARV16 Fab binding to immobilized MARV GPΔMuc by BLI, which revealed tight binding characterized by single digit nanomolar affinity (K_D_= 1.35 nM) **(Figure 3A; Table S1)**. Furthermore, MARV16 bound more strongly to MARV GPΔMuc than MR78 and MR191 in both IgG and Fab formats **(Figure 3B-C)**. Given that MARV enters target cells through fusion with the endosomal membrane induced by the low pH of late endosomes (*44–46*), we assessed the influence of pH on binding between MARV GPΔMuc and the three antibodies. We found that MARV16 bound comparably at all four pHs tested **(Figure 3D)** and that MR78 and MR191 binding was unaltered at pH 7.4, 6.5, and 5.5, and enhanced at pH 4.5 **(Figure S3)**, which we hypothesize may reflect increased accessibility of the RBS at lower pHs.

**Figure 3.**
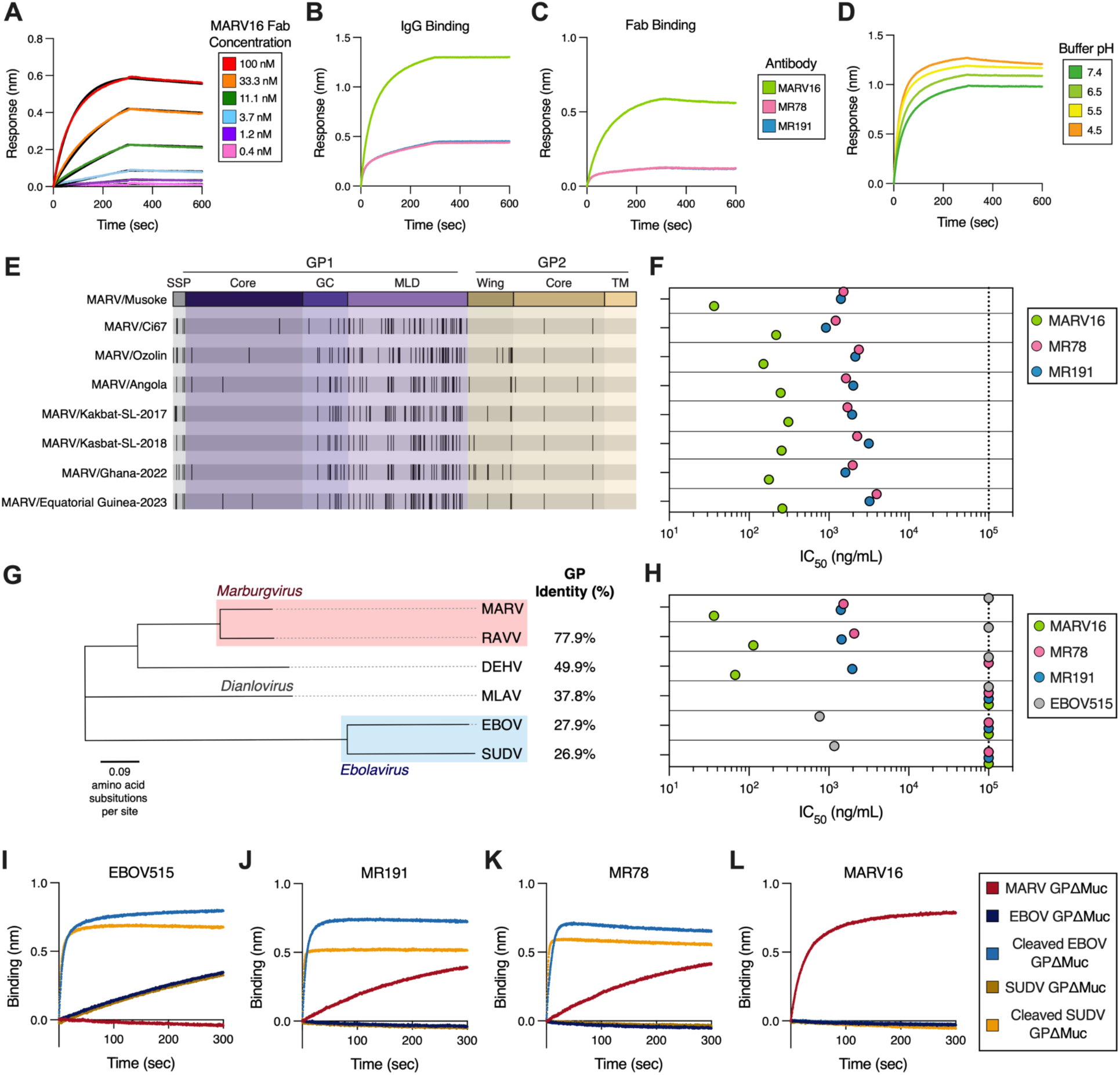
MARV16 potently neutralizes historical and contemporary MARV variants. **A)** Binding affinity determination of the MARV16 Fab to immobilized MARV GPΔMuc ectodomain as measured by BLI. Data presented are from one biological replicate and are representative of data from two biological replicates using distinct batches of protein. **B-C)** Comparison of binding of the MARV16, MR78, and MR191 (B) IgGs or (C) Fabs to immobilized MARV GPΔMuc ectodomain as assessed by BLI. Data presented are from one biological replicate and are representative of data from two biological replicates using distinct batches of protein. **D)** Comparison the binding of the MARV16 IgG to immobilized MARV GPΔMuc ectodomain at variable pHs using BLI. Data presented are from one biological replicate and are representative of data from two biological replicates using distinct batches of protein. **E)** Schematic highlighting the position of GP mutations (black vertical lines) in the MARV variants included in this study relative to the MARV/Musoke GP. **F)** Neutralization potency of MARV16, MR78, and MR191 against VSV pseudotyped with the indicated MARV GP. Data presented are averaged IC_50_ values obtained from at least two biological replicates conducted in technical triplicate using distinct batches of IgG and pseudoviruses. **G)** Phylogenetic tree constructed using the amino acid sequences of the filovirus GPs included in this study. The amino acid sequence identity of each GP compared to MARV/Musoke GP is shown to the right. **H)** Neutralization potency of MARV16, MR78, MR191, and EBOV515 against VSV pseudotyped with the indicated filovirus GP. **I-L)** Binding of the MARV GPΔMuc, EBOV GPΔMuc, SUDV GPΔMuc, thermolysin-cleaved EBOV GPΔMuc or thermolysin-cleaved SUDV GPΔMuc ectodomains to immobilized EBOV515 (I), MR191 (J), MR78 (K), or MARV16 (L) IgG as measured by BLI. Data presented reflect one biological replicate and are representative of two biological replicates using distinct batches of proteins.

We next evaluated the neutralization breadth of MARV16 against seven historical and contemporary MARV isolates using VSV pseudotyped with the corresponding MARV GPs which differ from that of the vaccine strain (MARV/Musoke) by 6.3 to 8.7% at the amino acid level **(Figure 3E)** with most substitutions mapping to the glycan cap or mucin-like domain. MARV16 potently neutralized all 7 vaccine-mismatched MARV isolates with an IC_50_ ranging from 151 to 310 ng/mL **(Figure 3F; Figure S4)** and markedly outperformed MR78 and MR191 against all of these MARV isolates when assessed side-by-side.

Evaluation of MARV16-mediated neutralization breadth across the *Filoviridae* family revealed that MARV16 potently inhibited RAVV and DEHV VSV pseudoviruses, but not Měnglà virus (MLAV), EBOV, or SUDV VSV **(Figure 3G-H; Figure S5)**. Previous studies showed that RBS-directed antibodies, including MR78 and MR191, recognize epitopes shared among all filovirus GPs but fail to neutralize *Ebolaviruses* due to masking mediated by the glycan cap (*15*, *35*). To determine whether MARV16 similarly recognizes a cryptic pan-filovirus epitope, we assessed whether the IgG binds the uncleaved and thermolysin-cleaved forms of the EBOV and SUDV GPΔMuc (i.e. removing the glycan cap) (*15*, *47*) and compared it to EBOV515, MR78 and MR191 using BLI. The *Ebolavirus* GP2-directed antibody EBOV515 bound uncleaved and cleaved EBOV and SUDV GPΔMuc, but not MARV GPΔMuc **(Figure 3I)**. MR78 and MR191 bound MARV GPΔMuc as well as the cleaved EBOV and SUDV GPΔMuc, but not the uncleaved EBOV and SUDV GPΔMuc **(Figure 3J-K)**. MARV16 bound the MARV GPΔMuc but not EBOV and SUDV GPΔMuc, irrespective of their cleavage, indicating that MARV16 does not cross-react with GPs from the *Filovirus* genus, most likely due to their extensive genetic divergence **(Figure 3G,L)**.

### Structural basis of MARV16-mediated neutralization

To understand the molecular basis of MARV16-mediated potent MARV neutralization, we characterized the MARV GPΔMuc ectodomain bound to the MARV16 Fab using cryogenic electron microscopy and determined a structure at 2.6 Å resolution **(Figure S6; Table S2)**. MARV16 recognizes a quaternary epitope spanning GP1 and GP2 **(Figure 4A-B)** which consists of residues 55-56, 58, 87-88, 90, 120, and 180 of GP1 and residues 505-507, 509-510, 512-513, 515-517, 550-552, 554, and 557 of GP2. An average surface area of 1,100 Å^2^ is buried at the interface between the epitope and the paratope upon binding with the majority of contacts with MARV GP contributed by the heavy chain. GP2 accounts for ∼75% of the epitope buried surface area and is recognized by all three complementarity-determining regions (CDRs) of the MARV16 heavy chain. CDRH3 residues recognize MARV GP2 through hydrogen bonds, salts bridges, and van der Waals interactions, including CDRH3 R99 and D105 forming salt bridges with GP2 E515 and K550, respectively, hydrogen bonding of the side chain nitrogen and backbone carbonyl oxygen of CDRH3 N101 with the side chains of GP2 E515 and N551, respectively, and hydrogen bonding of the CDRH3 W102 indol with GP2 N554 **(Figure 4C)**. CDRH1 and CDRH2 form extensive interactions with GP2, such as hydrogen bonding between the CDRH2 S53 and S54 side chains and GP2 D513, CDRH2 Y57 and the backbone amide and carbonyl oxygen of GP2 R517, the CDRH2 Y59 and GP2 R517 side chains. The CDRH1 S31 side chain contacts the GP2 G506 and E509 backbone carbonyl oxygens and CDRH1 T33 interacts with the GP2 E515 side chain. The MARV16 light chain additionally interacts with GP2 primarily through CDRL3 involving D93 salt bridged to GP2 R517 and hydrogen-bonding of the S91 and Y92 backbone carbonyls with the GP2 K550 side chain **(Figure 4D)**. MARV GP1 recognition is primarily mediated through MARV16 CDRH2 with the S54 and S56 backbone carbonyls respectively hydrogen-bonded to the GP1 K90 and K120 side chains and hydrogen-bonding of the CDRH2 Y57 and GP1 E87 side chains **(Figure 4E)**. These extensive contacts explain the strong MARV16 binding affinity and the conservation of interface residues among MARV isolates, RAVV, and DEHV rationalizes the cross-reactivity of this antibody **(Figure 3A,E-H).** Multiple epitope residue substitutions explain the lack of EBOV and SUDV VSV neutralization mediated by MARV16 **(Figure 3G-L)**.

**Figure 4.**
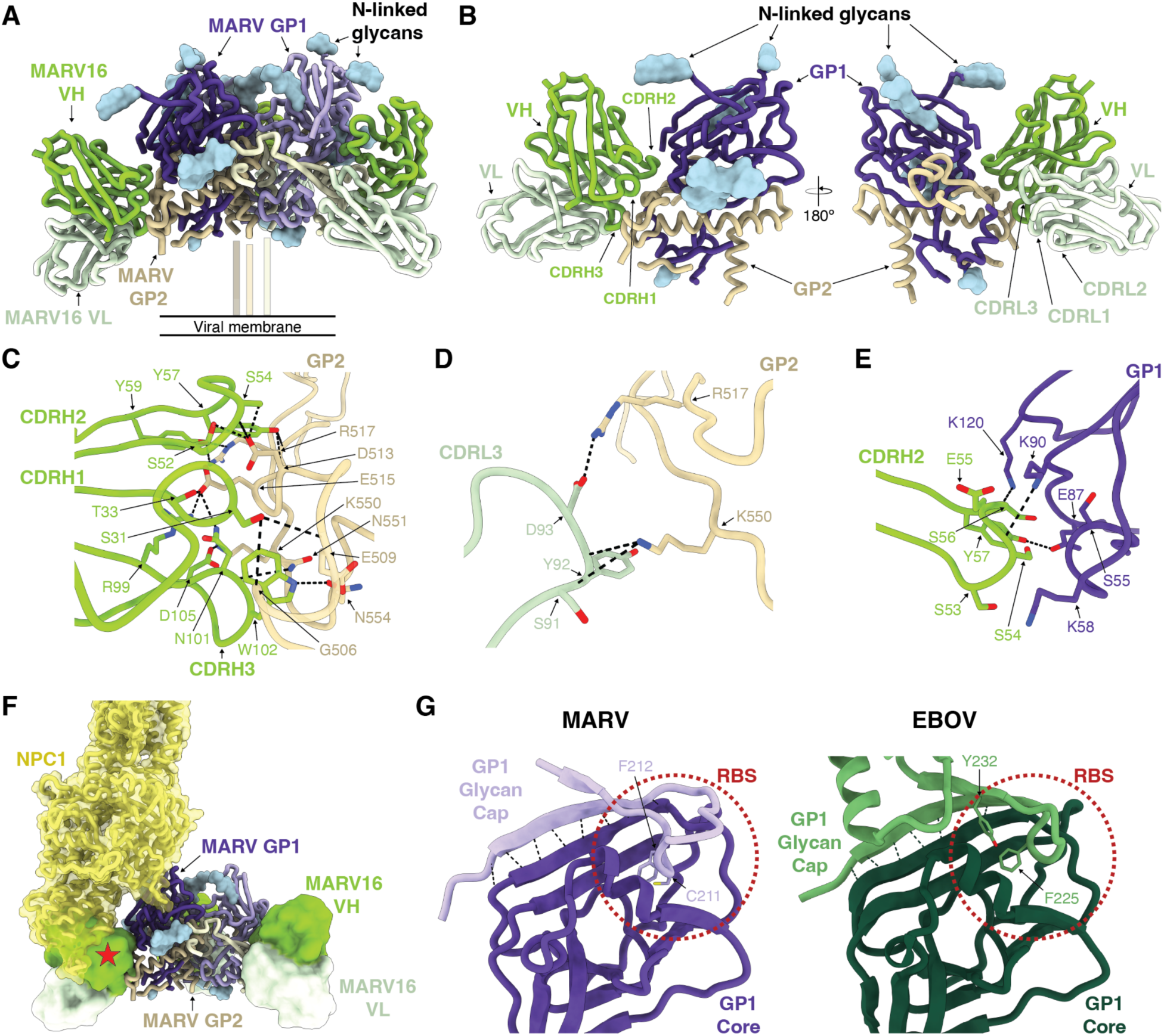
Molecular basis of MARV16-mediated MARV neutralization. **A)** Ribbon diagram of the cryo-EM structure of the MARV GPΔMuc ectodomain in complex with three MARV16 Fabs. Only the Fab variable domains were modeled into the density. MARV GP1 is shown in shades of purple, MARV GP2 is shown in shades of gold, MARV16 VH is shown in green, and MARV16 VL is shown in light green. N-linked glycan are rendered as blue surfaces. **B)** Ribbon diagram of a single MARV GP protomer in complex with one MARV16 Fab. **C-E)** Zoomed-in view of selected interactions between MARV16 CDRH1-3 and MARV GP2 (C), MARV CDRL3 and MARV GP2 (D), or MARV16 CDRH2 and MARV GP1 (E). Hydrogen bonds and salt bridges are denoted with black dashed lines. **F)** Binding modes of the NPC1 (yellow) and MARV16 (green/light green) to MARV GP. The position of NPC1 was determined by superimposing the EBOV GP-NPC1 model (PBD: 5JNX) with our MARV GP-MARV16 Fab model. The EBOV GP and the MARV GP glycan cap (residues 191-219) are not shown for clarity. The red star denotes the steric clash. **G)** Ribbon diagram of MARV and EBOV GP1 (PDB: 3CSY). The MARV GP1 core and glycan cap are shown in dark and light purple, respectively. The EBOV GP1 core and glycan cap are shown in dark and light green, respectively. Hydrogen bonds between the glycan cap and core are indicated with black dashed lines and glycan cap residues inserted in the receptor-binding site (RBS) are shown in stick representation and labeled. The position of the experimentally determined RBS for the EBOV GP and that of the putative RBS for the MARV GP are indicated with red dashed circles.

In the previously determined RAVV GP-MR191 (PBD: 6BP2) structure, the GP2 wing partially obstructs the MARV16 epitope (*16*). In our structure, the GP2 wing is largely disordered with the resolved wing residues shifting by up to 18 Å from their position in the RAVV GP-MR191 structure to accommodate binding of the MARV16 Fab **(Figure S7)**. These data indicate the wing is flexible and does not completely shield GP2 from neutralizing antibodies. Compared to structures of previously characterized anti-ebolavirus GP antibodies, MARV16 shares a similar binding mode to the EBOV GP-directed neutralizing antibody ADI-15946 **(Figure S8)**, which has been suggested to neutralize EBOV by tethering GP1 and GP2 in the prefusion conformation (*48*). Our structural data suggests that MARV16 locks the MARV GP1 and GP2 interface through contacts with residues that are rearranged during fusogenic conformational changes leading to membrane fusion (*21*). Furthermore, comparison with the NPC1-bound EBOV GP structure (*17*, *49*) indicates that MARV16 interferes with receptor binding as the heavy chain variable domain would sterically clash with the NPC1 N-terminal domain **(Figure 4F),** as is also the case for ADI-15946.

### Resolving the MARV GP glycan cap

The discovery of neutralizing antibodies targeting the MARV GP RBS from MARV-infected individuals suggested that the glycan cap might not be ordered and therefore does not shield the RBS (*35*, *36*). Our cryo-EM map resolves density near the RBS that corresponds to residues 191-219 of the GP1 glycan cap, consistent with partial shielding of the RBS by the glycan cap in a way reminiscent of that observed for EBOV GP **(Figure S9)**. Indeed, the EBOV GP glycan cap is interacting with the GP1 core via β-strand augmentation and insertion of F225 and Y232 into the RBS (*18*). Our structure reveals that the MARV GP architecture is strikingly similar to that of EBOV GP, sharing the β-strand augmentation and insertion of residues C211 and F212 into the MARV RBS **(Figure 4G)**. This difference may cause the MARV glycan cap to be more mobile and easily displaced than the EBOV glycan cap, explaining why MR78 and MR191 neutralize MARV, but not EBOV.

### Formulation of MARV neutralizing antibody cocktails

Antibody cocktails composed of two or more monoclonal antibodies targeting non-overlapping neutralizing epitopes are frequently used as antiviral therapeutics as they display greater resiliency to viral evolution than single monoclonal antibodies (*50*). As MARV16 binds to a distinct epitope on MARV GP than the RBS-directed MR78 and MR191, we reasoned that MARV16 and MR78 or MR191 can likely bind the MARV GP simultaneously. To examine this, we performed a competitive binding assay with MARV16 and MR78 or MR191 and observed that MARV GPΔMuc could bind MR78 or MR191 after binding MARV16 **(Figure 5A)**. Furthermore, we found that three MARV16 Fabs and three MR78 or MR191 Fabs could simultaneously bind to the prefusion MARV GPΔMuc ectodomain trimer, as visualized by electron microscopy of negatively-stained samples **(Figure 5B-C)**. These data indicate that MARV16 and an RBS-directed antibody can be used together in a therapeutic antibody cocktail to limit the risk of emergence of neutralization escape mutants.

**Figure 5.**
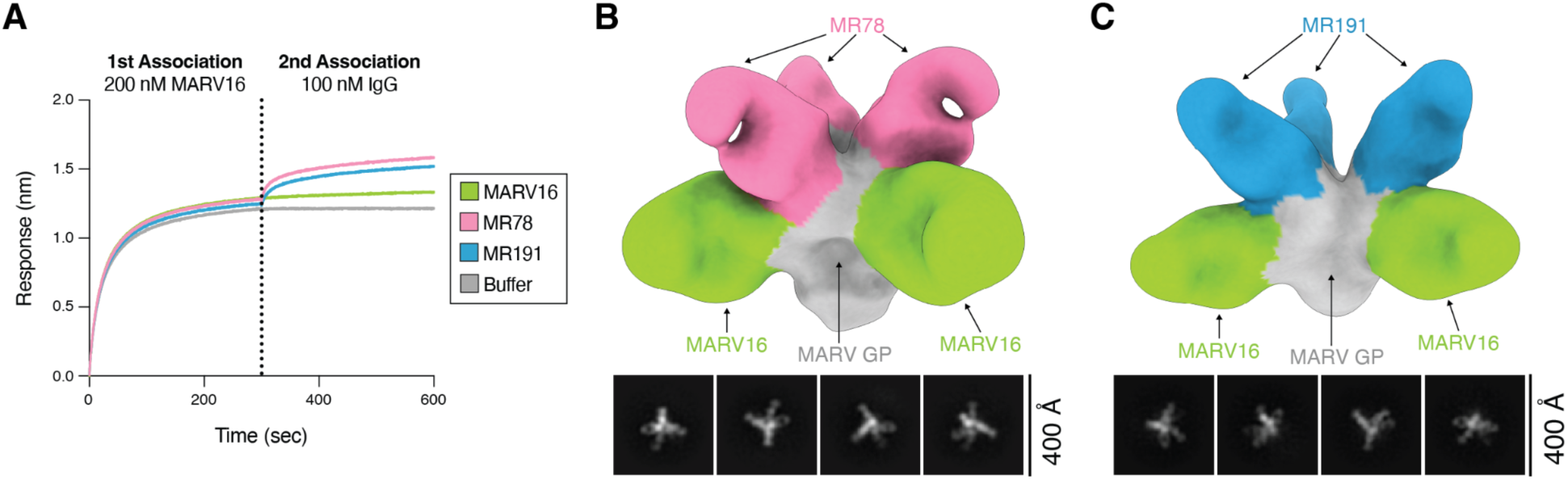
Formulation of a MARV monoclonal antibody cocktail. **A)** Competitive binding assay of MARV16, MR78, and MR191 IgG to the MARV16-bound MARV GPΔMuc ectodomain using BLI. Data presented are from one biological replicate and are representative of data from two biological replicates using distinct batches of protein. **B-C)** Representative 2D classes and 3D reconstruction of negatively stained MARV GPΔMuc ectodomain bound to MR78 and MARV16 Fabs (B) or MR191 and MARV16 Fabs (C). The position of the MR78 (pink) or MR191 (blue) Fabs were determined by superimposing the RAVV GP-MR78 Fab (PDB: 5UQY) or RAVV GP-MR191 Fab (PDB: 6BP2) structures with our MARV GP-MARV16 Fab structure.

## Discussion

Identification of stabilizing mutations is a key goal of vaccine design as it can markedly improve the immunogenicity of viral antigens by preferentially eliciting antibodies directed towards the desired conformation of a glycoprotein, as demonstrated by the success of the SARS-CoV-2 S and RSV F vaccines which incorporate prefusion-stabilizing mutations (*28–32*, *51*). The stabilizing mutations identified here improve both expression and thermostability of MARV GPΔMuc and will likely improve the immunogenicity of MARV vaccines in development. Furthermore, we used ProteinMPNN (*43*) to identify stabilizing mutations in the MARV GPΔMuc similar to approaches recently used for the Langya virus G and Epstein-Barr virus gB proteins (*52*, *53*), heralding a new era of machine learning-enabled vaccine design.

Prior studies suggested that the MARV GP equator and base are shielded from neutralizing antibodies by the GP1 mucin-like domain and the GP2 wing (*15*, *16*) given that all previously characterized MARV GP neutralizing antibodies solely target the RBS (*35*, *36*). As MARV16 binds to an epitope spanning GP1 and GP2, our data show that the mucin-like domain and wing do not fully shield GP2 but instead demonstrate that the wing is conformationally flexible, thereby enabling antibody binding. As a result, we anticipate that future antibody discovery campaigns will identify neutralizing antibodies targeting multiple different GP2 epitopes. MARV16 recognizes a conserved epitope, neutralizing filoviruses as distantly related as DEHV, which only shares 49.9% GP amino acid identity with MARV (*54*). Several *Ebolavirus* GP-directed antibodies that neutralize EBOV, SUDV, and Bundibugyo ebolavirus, but not MARV, including ADI-15946, EBOV-515, and EBOV-520, recognize similar epitopes to MARV16 (*48*, *55–57*), indicating this epitope is a prime target for broad genus-specific neutralization. Therapeutics or vaccines targeting this MARV GP antigenic site will thus likely provide robust protection against pre-emergent MARV variants and MARV-related filoviruses, similar to those developed for EBOV (*58*).

For *Ebolaviruses*, a structured glycan cap blocks access to the RBS until GP cleavage mediated by cathepsin B or L in the endosomes (*18*, *19*, *44*, *59*, *60*), thereby limiting the elicitation of and potency of RBS-directed antibodies (*15*, *35*). In contrast, the glycan cap had not been visualized for MARV GP and RBS-directed antibodies mediate MARV neutralization, suggesting that the MARV GP RBS is more exposed than that of *Ebolavirus* GPs (*15*, *16*, *35*, *36*). Although our structure reveals that the MARV glycan cap shields the RBS similarly to the glycan cap of *Ebolaviruses*, increased conformational heterogeneity or looser tethering of the MARV GP glycan cap to the RBS might enable easier displacement, explaining the greater neutralizing activity of MR78 and MR191 against MARV, relative to *Ebolaviruses*. Accordingly, we observed an increase in binding for the RBS-directed antibodies MR78 and MR191 to the MARV GPΔMuc ectodomain in acidic conditions. These data suggest the glycan cap is less likely to mask the RBS in the acidic conditions of late endosomes.

Therapeutic antibody cocktails consisting of multiple antibodies targeting unique epitopes on an antigen are favored for viral pathogens as the targeted viral protein would typically need to accumulate multiple mutations to evade all the antibodies in the cocktail (*50*). All prior neutralizing antibodies identified against the MARV GP target the RBS, which limited the development of an antibody cocktail for MARV (*35–38*). We identified a best-in-class antibody targeting a previously unknown MARV GP antigenic site and demonstrated that MARV16 can bind to the MARV GP concurrently with RBS-directed antibodies. These data indicate a therapeutic antibody cocktail against MARV, similar to ZMapp for EBOV (*61*), can be developed.

In summary, our results will inform both vaccine and therapeutic development against MARV, providing improved treatment and prevention options for future MARV outbreaks.

## Acknowledgments

This study was supported by the National Institute of Allergy and Infectious Diseases (DP1AI158186 and 75N93022C00036 to D.V.), an Investigators in the Pathogenesis of Infectious Disease Awards from the Burroughs Wellcome Fund (D.V.), the University of Washington Arnold and Mabel Beckman cryo-EM center and the National Institute of Health grant S10OD032290 (to DV). DV is an Investigator of the Howard Hughes Medical Institute and the Hans Neurath Endowed Chair in Biochemistry at the University of Washington.

## Author contributions

AA, LP, DC, FB and DV conceived the project. AA, JTB, and CS recombinantly expressed glycoproteins. AA designed MARV GP constructs and evaluated their antigenicity and stability. LP and AD performed immunization, mAb isolation and ELISAs. KC and AB produced the recombinant antibodies. MG and RC carried out authentic virus neutralization assays. AA conducted biolayer interferometry binding experiments and pseudovirus neutralization assays. AA carried out negative stain and cryo-EM sample preparation, data collection, and processing with help from YJP and MM. AA and DV built and refined the structures. AA and DV wrote the manuscript with input from all authors. AA, LP, DC, FB and DV analyzed the data.

## Methods

### Cells

HEK-293T and Vero E6 cells were grown in DMEM supplemented with 10% FBS and 1% PenStrep at 37°C and 5% CO_2_. Expi293 were grown in Expi293 media at 37°C and 8% CO_2_ rotating at 130 RPM.

### In vivo animal studies

ATX-GK and ATX-GL female mice, 6-7 weeks old, were obtained from Alloy Therapeutics Inc. and housed for the immunization experiment at the Institute for Research in Biomedicine, Bellinzona, Switzerland. All animal experiments were performed in accordance with the Swiss Federal Veterinary Office guidelines and authorized by the Cantonal Veterinary (approval no. 35554 TI-39/2023/2023). Animals were supervised by a licensed veterinarian and proper steps were taken to ensure the welfare and minimize the suffering of all animals in the conducted studies. Animals were housed in ventilated cages in a 12 h light/dark cycle, with free access to water and standard sterilized chow.

### Constructs

The construct encoding the MARV GPΔMuc ectodomain (residues 1-256 and 426-637) with a C-terminal 8x His tag was codon optimized, synthesized, and inserted into pcDNA3.1(+) by Genscript. Mutations neighboring the furin cleavage site (W439A, F445G, and F447N) and the H589I stabilizing mutation were introduced using In-Fusion Cloning with overlapping mutagenesis primers. The HR1_c_ stabilizing mutations (T582P and F583V) were additionally introduced through In-Fusion Cloning using overlapping mutagenesis primers. Constructs encoding the EBOV GPΔMuc (residues 1-312 and 463-637) and SUDV GPΔMuc (residues 1-313 and 474-637) both with a C-terminal T4 foldon and 8x His tag were codon optimized, synthesized, and inserted into pcDNA3.4 by Genscript. The construct encoding the full-length MARV/Musoke GP (GenBank Accession Number: NC_001608) with a C-terminal FLAG tag was codon optimized, synthesized, and inserted into pcDNA3.1(+) by Genscript. Constructs encoding the full-length MARV/Ci67 (GenBank Accession Number: EF446132), MARV/Ozolin (GenBank Accession Number: AY358025), MARV/Angola (GenBank Accession Number: KY047763), MARV/Kakbat-SL-2017 (GenBank Accession Number: MN258361), MARV/Kasbat-SL-2018 (GenBank Accession Number: MN187403), MARV/Ghana-2022 (GenBank Accession Number: OQ672470), and MARV/Equatorial Guinea-2023 (HS415030) were codon optimized, synthesized, and inserted into pHDM by Genscript. Constructs encoding the MR78, MR191, and EBOV515 heavy and light chains were codon optimized, synthesized, and inserted into pcDNA3.1(+) by Genscript.

### Recombinant protein expression and purification

To produce the MARV GPΔMuc, EBOV GPΔMuc, and SUDV GPΔMuc ectodomains, Expi293 cells were grown to a density of 3 x 10^6^ cells/mL and then transfected with constructs encoding MARV GPΔMuc and furin at a 3:1 mass ratio using the Expifectamine293 transfection kit following the manufacturer’s instructions. Five days after transfection, the supernatant was collected, clarified by centrifugation, and incubated with Ni Sepharose Excel resin (Cytiva) for 1 hour at room temperature. The resin was then collected in a gravity column and washed with wash buffer (25 mM sodium phosphate pH 8.0, 300 mM NaCl, and 50 mM imidazole or 100 mM Tris pH 8.0, 300 mM NaCl, and 40 mM imidazole). The proteins were then eluted using an elution buffer containing 25 mM sodium phosphate, 300 mM NaCl, 500 mM imidazole, pH 8.0 or 100mM Tris, 300 mM NaCl, 300 mM imidazole and further purified into TBS (20 mM Tris pH 7.4 and 100 mM NaCl, or 50 mM Tris pH 7.4 and 150 mM NaCl) by size-exclusion chromatography using a Superose 6 Increase 10/300 GL column. The purified proteins were concentrated using a 100-kDa Amicon centrifugal filter, flash frozen, and stored at -80°C until use.

MR78, MR191, and EBOV515 monoclonal antibodies were produced by transfecting Expi293 cells grown to a density of 3 x 10^6^ cells/mL with the heavy chain and light chain constructs supplied at a 1:1 mass ratio using the Expifectamine293 transfection kit. Four to 5 days after transfection, the supernatant was collected, clarified by centrifugation, and flowed over a Protein A column. The column was then washed with at least ten column volumes of wash buffer containing 20 mM sodium phosphate, pH 8.0. The eluted antibodies were then exchanged into TBS and concentrated using a 100-kDa Amicon centrifugal filter.

### Biotinylation of MARV GPΔMuc

The MARV GPΔMuc was produced and purified as described above. Following elution from the Ni Sepharose Excel resin, the GP was exchanged into PBS (137 mM NaCl, 2.7 mM KCl, 10 mM Na_2_HPO_4_, 1.8 mM KH_2_PO_4_, pH 7.4) and concentrated to 1 mg/mL using a 100-kDa Amicon centrifugal filter. The MARV GPΔMuc was biotinylated using the EZ-Link Sulfo-NHS-SS-Biotinylation Kit (ThermoFisher) using a 40-fold molar excess of Sulfo-NHS-SS-biotin and incubating the reaction at room temperature for 30 minutes. The biotinylated MARV GPΔMuc was then purified into TBS by size-exclusion chromatography using a Superose 6 Increase 10/300 GL column. The purified protein was then concentrated using a 100-kDa Amicon centrifugal filter, flash frozen, and stored at -80°C until use.

### Cleavage of EBOV and SUDV GPΔMuc

The EBOV and SUDV GPΔMuc ectodomains were produced and purified as described above. Following elution from the Ni Sepharose Excel resin, the GP was exchanged into TBS and concentrated to 1 mg/mL using a 100-kDa Amicon centrifugal filter. Thermolysin, resuspended in TBS, was added to a final concentration of 0.2 mg/mL. The reaction was incubated at 37°C for 1 hour after which phosphoramidon was added to a final concentration of 500 µM to stop the thermolysin. The cleaved GP was purified into TBS by size-exclusion chromatography using a Superdex 200 Increase 10/300 GL column.

### Generation of Fabs

To generate Fabs from the purified monoclonal antibodies, LysC, resuspended in TBS, was added to 1 mg of MARV16, MR78, or MR191 IgG at a 1:4,000 to 1:8,000 mass ratio and incubated at 37°C overnight. The following day, MabSelect Resin was added to the digested IgG solution and incubated for 1 hour at room temperature. The flowthrough was collected and run over a Superdex 75 Increase 10/300 GL column into TBS. Fractions containing the Fab were pooled and concentrated using a 30-kDa Amicon centrifugal filter.

### MARV GPΔMuc immunizations and antibody discovery from Alloy Mice

Pre-immune serum was obtained from each mouse a week before immunization. ATX mice were immunized with recombinant MARV GPΔMuc diluted (1:1) in Magic Mouse adjuvant (Cat#: CDN-A001E; CD Creative Diagnostics) and injected subcutaneously and intraperitoneally. On day 0, mice received prime immunization with 20ug of MARV GPΔMuc and were boosted on day 13 and day 64 with the same amount of antigen. On day 70, the mice were sacrificed and peripheral blood, spleen and lymph nodes (LN) were collected and cells freshly isolated. B cells from either freshly isolated or frozen splenocytes were enriched by positive selection using mouse CD19 microbeads and LS columns (Miltenyi) and subsequently stained with mouse anti-IgM, anti-IgD, anti-IgA and biotinylated MARV GPΔMuc labeled with both streptavidin-Alexa-Fluor 488 and streptavidin-Alexa-Fluor 647 (Life Technologies). MARV GPΔMuc -specific IgG+ memory B cells were sorted by flow cytometry via gating out IgM/IgD/IgA-positive B cells and positively baiting B cells with dual-labeled (Alexa-Fluor 488 and Alexa-Fluor 647) antigen, using SH800SFP cell sorter (Sony). Sorted IgG+ memory B cells were seeded at clonal dilution in 384-well plates on a monolayer of feeder mesenchymal cells in the presence of B cell survival factors. Clones positive for antigen binding were then isolated and the cDNA synthesized. mAb VH and VL sequences were obtained by reverse transcription PCR (RT-PCR), The V, D and J genes of the IgH DNA sequences were identified using the IMGT database as a reference (*62*). mAbs were then produced recombinantly as human IgG1 (IgG1m3 allotype) in ExpiCHO cells, transiently transfected with heavy and light expression vectors as previously described (*63*).

### Enzyme-linked immunosorbent assay (ELISA)

The MARV GPΔMuc ectodomain was diluted to 0.003 mg/mL in TBS, added to Maxisorp 384-well plates (ThermoFisher), and incubated overnight at room temperature. The following day, the plates were slapped dry and blocked with Blocker Casein for 1 hour at 37°C. The plates were slapped dry again and the monoclonal antibodies were diluted to a starting concentration of 0.1 mg/mL in TBS with 0.1% Tween 20 (TBST). The antibodies were serially diluted 1:3 in TBST thereafter, added to plates, and incubated at 37°C for 1 hour. The plates were slapped dry and washed four times with TBST after which a goat anti-human IgG (H+L) HRP conjugated antibody (ThermoFisher) diluted 1:5,000 in TBST was added to each well. The plates were incubated for 1 hour at 37°C, slapped dry, and washed four times with TBST. SureBlue Reserve TMB 1-Component Microwell Peroxidase Substrate (SeraCare) was added to each well and developed for 90 seconds at room temperature. An equal volume of 1N HCl was added to each well to quench the reaction after which the absorbance at 450 nm was measured using a BioTek Synergy Neo2 plate reader. The resulting data were analyzed using GraphPad Prism 10 using a four parameter logistic curve to determine the ED_50_ for each antibody. Two biological replicates performed in technical duplicate were performed using two distinct batches of protein.

### Pseudotyped virus production

Vesicular stomatitis virus (VSV) pseudotyped with the full length MARV, RAVV, DEHV, MLAV, EBOV, or SUDV GP was produced as previously described (*51*, *64–66*). In brief, HEK-293T cells were split into 10 cm poly-lysine coated dishes and grown overnight at 37°C and 5% CO_2_ until they reach approximately 90-95% confluency. The cells were washed once with DMEM and left in DMEM supplemented with 10% FBS. The cells were transfected with 16-24 µg of full length GP construct using Lipofectamine 2000 following the manufacturer’s recommendations after which the cells were incubated for 20-24 hours at 37°C and 5% CO_2_. The cells were then washed three times with DMEM, infected with VSVΔG/luc, and incubated 37°C and 5% CO_2_. After two hours, the cells were washed five times with DMEM and left in DMEM supplemented with an anti-VSV-G antibody (I1-mouse hybridoma supernatant diluted 1:25 for CRL-2700, ATCC) for 20-24 hours at 37°C and 5% CO_2_. Following this incubation, the supernatant was collected, clarified by centrifugation, filtered using a 0.45 µM filter, and concentrated with a 100-kDa centrifugal filter (Amicon). The resulting pseudovirus was stored at -80°C until use.

### Pseudovirus neutralization assay

Neutralization assays were performed as previously described (*51*, *64–66*). Briefly, Vero E6 cells were split into white walled, clear bottom 96-well plates at a density of 18,000 cells per well. The cells were grown overnight at 37°C and 5% CO_2_ until they reached approximately 80-90% confluency. The monoclonal antibodies were diluted in DMEM to a starting concentration of 200 µg/mL and serially diluted 1:3 in DMEM thereafter. VSV pseudotyped with the GP was diluted 1:5 to 1:250 in DMEM after which an equal volume of diluted pseudovirus was added to the diluted monoclonal antibody. The pseudovirus-antibody mixture was incubated at room temperature for 30 minutes. Following this incubation, growth media was removed from the Vero E6 cells and the pseudovirus-antibody mixture was added to cells. The cells were incubated for 2 hours at 37°C and 5% CO_2_ after which an equal volume of DMEM supplemented with 20% FBS and 2% PS was added to each well and the cells were incubated for another 20-24 hours at 37°C and 5% CO_2_. An equal volume of ONE-Glo EX was added to each well and the cells were incubated 37°C for 5 minutes with constant shaking. The luminescence values from each well were measured using a BioTek Synergy Neo2 plate reader.

Data were normalized using GraphPad Prism 10 using the relative light unit (RLU) values obtained from uninfected cells to define 100% neutralization and the RLU values obtained from cells infected with pseudovirus in the absence of antibody to define 0% neutralization. ED_50_ values were determined from the normalized data using an [inhibitor] vs. normalized response -variable slope model. At least two biological replicates using distinct batches of pseudoviruses and monoclonal antibodies were performed.

### Plaque reduction neutralization assay with authentic MARV

VeroE6 cells were split into 6-well plates at a density of 9 x 10^5^ cells per well and grown overnight in DMEM supplemented with 10% FCS at 37°C and 5% CO_2_. The following day, monoclonal antibodies were diluted in DMEM to a starting concentration of 100 µg/mL and serially diluted 1:4 in DMEM thereafter. Next, 100 µL of MARV diluted to 1,000 PFU/mL was added to 100 µL of the diluted antibodies and the virus-antibody mixture was incubated for 60 minutes at 37°C. Following this incubation, an additional 300 µL of DMEM was added to the virus-antibody mixture and 400 µL of this mixture was added to the VeroE6 cells. The cells were incubated with the virus-antibody mixture for 60 minutes at 37°C after which the mixture was removed and an overlay consisting of 1% Seakem agarose (Lonza) mixed 1:1 with 2x Eagle’s MEM (EMEM; Lonza) containing 4 mM L-glutamine, 2 mM sodium pyruvate, and 4% FCS was added. The cells were incubated for 7 days at 37°C with 5% CO_2_. The cells were stained with an overlay containing 1% Seakem agarose mixed 1:1 with EMEM containing 4 mM L-glutamine, 2 mM sodium pyruvate, 4% FCS, and 8% neutral red solution and incubated for 1 day at 37°C and 5% CO_2_ after which the number of plaques were counted. The percent infectivity for each well was determined by dividing the number of plaques in the well by the number of plaques counted in the well with 2.4 x 10^-5^ µg/mL of antibody. Two biological replicates with one to three technical replicates were conducted for each antibody and the PRNT_50_ values were determined from the averaged data from the two biological replicates using an [inhibitor] vs. normalized response - variable slope model in GraphPad Prism 10.

### Biolayer interferometry

Binding of the stabilized MARV GPΔMuc ectodomains to MR191 was assessed by first dipping pre-hydrated AHC2 biosensors into MR191 IgG diluted to 10 ng/µL in 10x kinetics buffer to a 1 nm shift. The MR191-coated biosensors were then dipped into MARV GPΔMuc diluted to 10 nM in 10x kinetics buffer for 300 s after which the biosensors were dipped into 10x kinetics buffer. All steps were conducted at 30°C. Data were baseline subtracted using Octet Data Analysis HT software v12.0 and visualized using GraphPad Prism 10.

To measure the affinity of the MARV16 Fab for the MARV GPΔMuc, biotinylated MARV GPΔMuc was diluted to a concentration of 10 ng/µL in 10x kinetics buffers and loaded onto pre-hydrated streptavidin biosensor to a 1 nm shift. The MARV GPΔMuc-coated biosensors were then dipped into MARV16 Fab diluted in 10x kinetics buffer at a starting concentration of 100 nM and serially diluted 1:3 thereafter for 300 s. The biosensors were then dipped into 10x kinetics buffer for an additional 300 s. All steps were conducted at 30°C. The resulting data were baseline subtracted and fit using Octet Data Analysis HT software v12.0 and visualized using GraphPad Prism 10. Affinity comparisons between the MARV16, MR78, and MR191 Fabs and IgGs were conducted similarly as described above. Following immobilization of the biotinylated MARV GPΔMuc on the streptavidin biosensors, the tips were dipped into 100 nM of Fab or IgG diluted in 10x buffer for 300 s after which the tips were dipped into 10x kinetics buffer for 300 s. All steps were conducted at 30°C. Data were baseline subtracted using Octet Data Analysis HT software v12.0 and visualized using GraphPad Prism 10.

Binding of MARV GPΔMuc to MARV16, MR78, or MR191 IgG at variable pHs were conducted by loading biotinylated MARV GPΔMuc diluted in 10x kinetics buffer, pH 7.4 onto streptavidin biosensors to a 1 nm shift. The loaded biosensors were then dipped into IgG diluted in 10x kinetics buffer at pH 7.4, 6.5, 5.5, or 4.5 for 300 s. The resulting data were baseline subtracted using Octet Data Analysis HT software v12.0 and visualized using GraphPad Prism 10.

Binding of the MARV GPΔMuc, EBOV GPΔMuc, cleaved EBOV GP, SUDV GPΔMuc, and SUDV GPΔMuc to MARV16, MR78, MR191, and EBOV515 IgG was assessed similarly as described above. IgG diluted to 10 ng/uL in 10x kinetics buffer was loaded on AHC2 biosensors to a 1 nm shift after which the loaded biosensors were dipped into GP diluted to approximately 10 nM in 10x kinetics buffer for 300 s. All steps were conducted at 30°C. Data were baseline subtracted using Octet Data Analysis HT software v12.0 and visualized using GraphPad Prism 10.

Competitive binding of MARV16 versus MR78 or MR191 to the MARV GPΔMuc was assessed by loading biotinylated MARV GPΔMuc diluted to 10 ng/µL onto pre-hydrated streptavidin biosensor after which the loaded biosensors were dipped into 200 nM of MARV16 IgG diluted 10x kinetics buffer for 300 s. The biosensors were then dipped into 100 nM of MR78, MR191, or MARV16 IgG diluted in 10x kinetics buffer or 10x kinetics buffer for 300 s and finally dipped into 10x kinetics buffer for 300 s. All steps were conducted at 30°C. The resulting data were baseline subtracted using Octet Data Analysis HT software v12.0 and visualized using GraphPad Prism 10.

### Cryo-EM sample preparation and data collection

To generate MARV GPΔMuc-MARV16 Fab complexes, 100 µg of MARV GPΔMuc and 150 µg MARV16 Fab were incubated together at 37°C for 30 minutes after which the complexes were added to a 100 kDa centrifugal filter, washed 5 times with TBS, and concentrated to 5.5 mg/mL. Immediately prior to freezing, CHAPSO was added to the MARV GPΔMuc-MARV16 Fab complexes to a final concentration of 5.45 mM. The complexes were added to freshly glow discharged 2.0/2.0 UltraFoil grids (200 mesh) (*67*) after which the grids were plunge frozen using a Vitrobot MarkIV (ThermoFisher) with a wait time of 10 s, a blot force of 0, and a blot time of 5 s at 100% humidity and 23°C. Data were acquired on a FEI Titan Krios transmission electron microscope operated at 300 kV and equipped with a Gatan K3 direct detector and Gatan Quantum GIF energy filter, operated in zero-loss mode with a slit width of 20 eV. Automated data acquisition was carried out using Leginon (*68*) at a nominal magnification of 105,000x with a pixel size of 0.843 Å, a defocus range between -0.4 to -3.0 µm, and a stage tilt of 0° or 25°. The dose rate was adjusted to 15 counts/pixel/s and each movie was acquired in 75 frames of 40s.

### Cryo-EM data processing

Movie frame alignment with a downsampled pixel size of 1.658 Å was carried out in WARP (*69*). Estimation of microscope CTF parameters, particle picking, and extraction (box size: 256 pixels^2^) was conducted using cryoSPARC. Reference-free 2D classification to select well-defined particle images was performed in cryoSPARC (*70*). Next, ab-initio 3D reconstruction and heterologous refinement to select well-defined particle classes were performed in cryoSPARC. 3D refinements were then conducted using non-uniform refinement (*71*) with per-particle defocus refinement in cryoSPARC (*71*). The particle images were then subjected to Bayesian polishing using Relion (*72*) during which the box size was adjusted to 512 pixels^2^ and the pixel size was adjusted to 0.829 Å. Another round of non-uniform refinement with global and per-particle defocus refinement was performed in cryoSPARC. Next, focused 3D classification was conducted using a mask over residues 171-219 using a tau factor of 10 in Relion (*73*, *74*). The particles with the best resolved local density were selected and subjected to local refinement in cryoSPARC using a mask over the MARV GP and Fab variable domains. Reported resolutions are based on the gold-standard Fourier shell correlation of 0.143 criterion and Fourier shell correlation curves were corrected for the effects of soft masking by high-resolution noise substitution (*75*, *76*).

### Model building and refinement

USCF ChimeraX (*77*) and Coot (*78*) were used to fit initial models of the MARV GP (PDB: 6BP2) and MARV16 Fab, which was predicted using AlphaFold2 (*79*), into the map. The model was then built and refined into the map using Coot, Rosetta (*80*, *81*), ISOLDE (*82*), and Phenix (*83*). Figures were generated using UCSF ChimeraX.

### DIfferential scanning fluorimetry

The original and stabilized MARV GPΔMuc ectodomains were diluted in TBS and mixed with Protein Thermal Shift Buffer and Dye (ThermoFisher) following the manufacturer’s recommendation such that the final concentration of protein in the reaction mix was 0.25 µg/mL. The reaction mix was added to a 96-well qPCR plate (ThermoFisher) and sealed with MicroAmp Optical Adhesive Film (ThermoFisher). The fluorescence intensity (λ_Excitation_: 470 ± 15 nm; λ_Emission_: 586 ± 10 nm) was measured from 25°C to 99°C using a QuantStudio 5 Real-Time PCR System (ThermoFisher). The data were analyzed and visualized using QuantStudio Design and Analysis Desktop Software v (ThermoFisher) and GraphPad Prism 10. Data are presented as the negative first derivative of fluorescence intensity with respect to temperature. The melting temperature was identified by taking the second derivative of the fluorescence intensity with respect to temperature and smoothing the resulting function across 4 neighbors points. Four biological replicates each with six technical replicates were performed using distinct batches of protein.

### Negative stain electron microscopy

Complexes of the MARV GPΔMuc-MARV16 Fab-MR78 Fab or MARV GPΔMuc-MARV16 Fab-MR191 Fab were generated as described above. Purified MARV GPΔMuc mutants or MARV GPΔMuc-Fab complexes were diluted to 0.01 mg/mL in TBS, added to freshly glow-discharged carbon-coated copper grids, and stained with 2% uranyl formate. Data were acquired with a 120kV FEI Tecnai G2 Spirit with a Gatan Ultrascan 4000 4k x 4k CCD camera at a nominal magnification of 67,000x using Leginon (*68*). The defocus ranged from -3.0 to -1.0 µm and the pixel size was 1.6 Å. Micrographs were then processed in cryoSPARC (*70*) using PatchCTF to estimate microscope CTF parameters and Blob picker to pick particles. Following particle extraction, reference-free 2D classification was performed to select well-defined particle images. Ab-initio 3D reconstruction was then performed with the selected particle images applying C3 symmetry. Figures were generated using UCSF ChimeraX (*77*).

## Data Availability

The cryo-EM maps and atomic coordinates will be deposited to the Electron Microscopy Data Bank (EMDB) and the PDB. Data generated in the study are available from the corresponding authors. Materials generated in this study can be available on request and may require a material transfer agreement.

---

**Figure S1.**
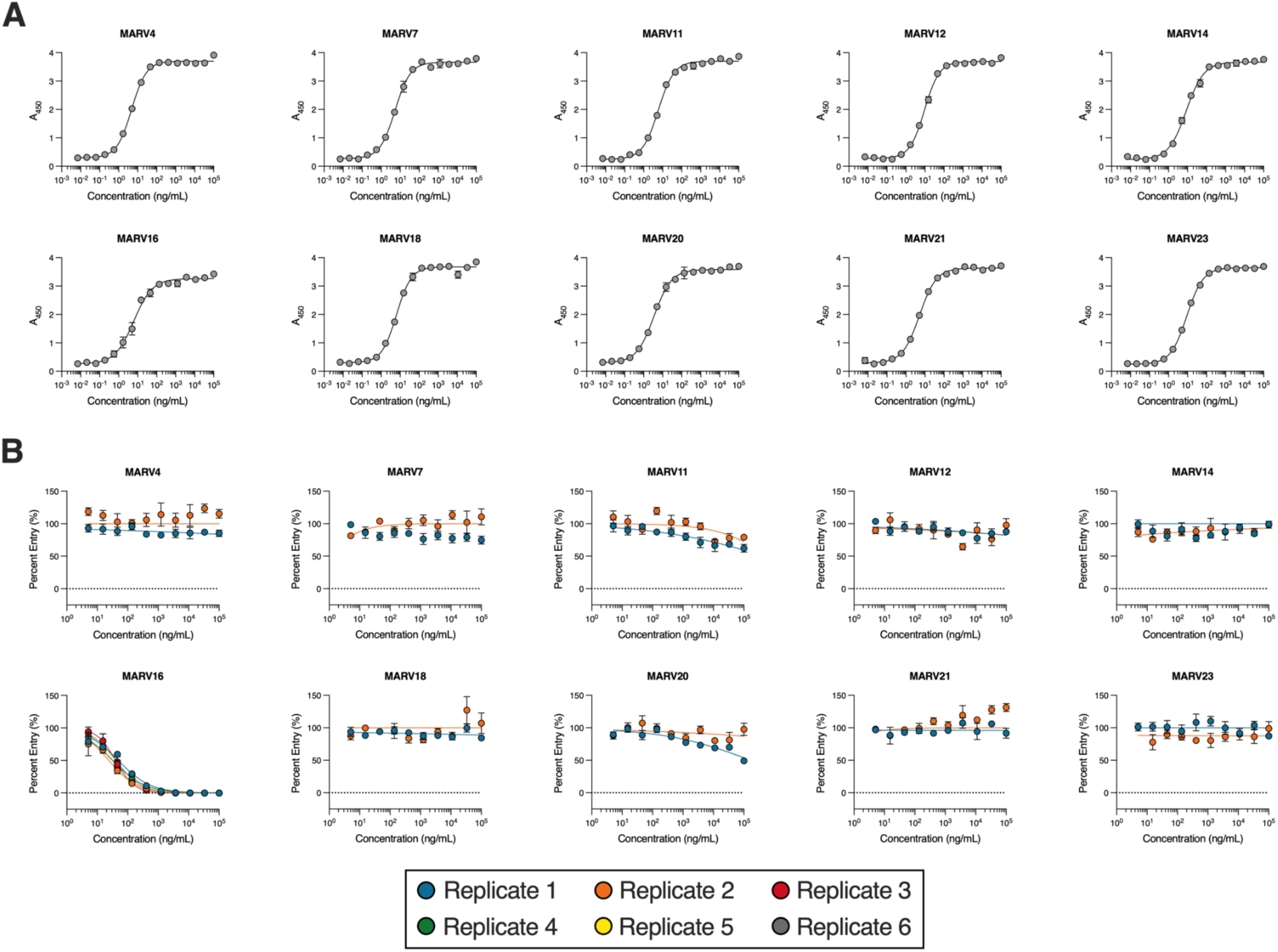
Dose-response curves for the ELISAs against the MARV GPΔMuc ectodomain (A) and neutralization assays (B) against VSV pseudotyped with MARV/Musoke GP for the 10 antibodies discovered from the immunization study using the ATX-GK mouse. Two biological replicates were performed for the ELISAs using distinct batches of proteins and antibodies. Two technical replicates were performed per biological replicate. Data presented are from one representative biological replicate and presented as mean ± standard error from the two technical replicates. Two to six biological replicates were performed for the neutralization assays using distinct batches of antibodies and pseudoviruses. Three technical replicates were performed per biological replicate. Data from all biological replicates are shown and presented as mean ± standard error from the three technical replicates.

**Figure S2.**
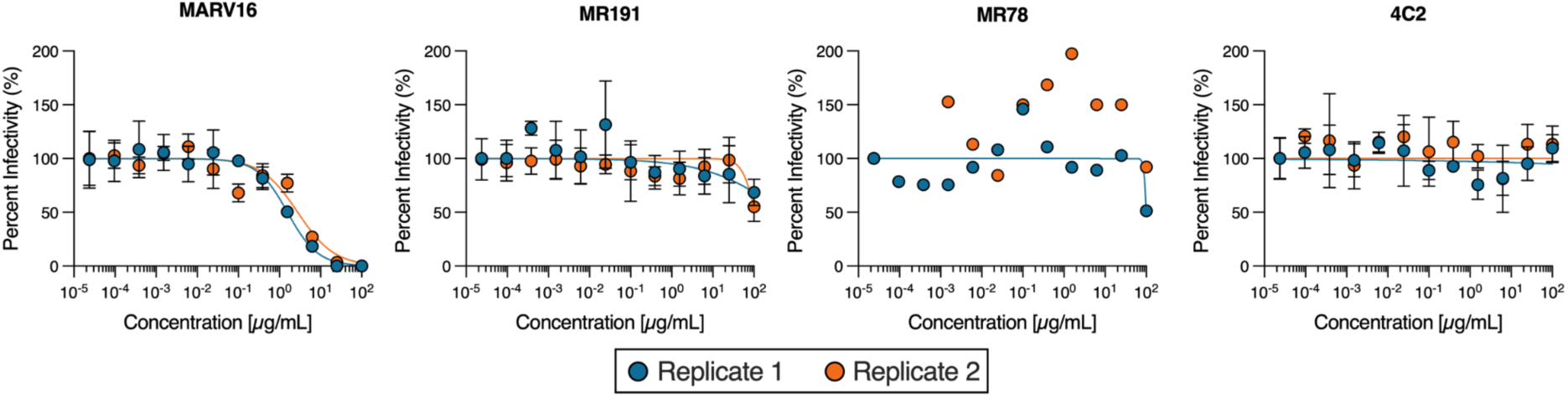
Dose-response curves for plaque reduction neutralization tests for MARV16, MR78, MR191, and 4C2, conducted using authentic MARV/Musoke. Two biological replicates were performed with one to three technical replicates using distinct batches of monoclonal antibodies. Data are shown as the mean ± standard error of the technical replicates.

**Figure S3.**
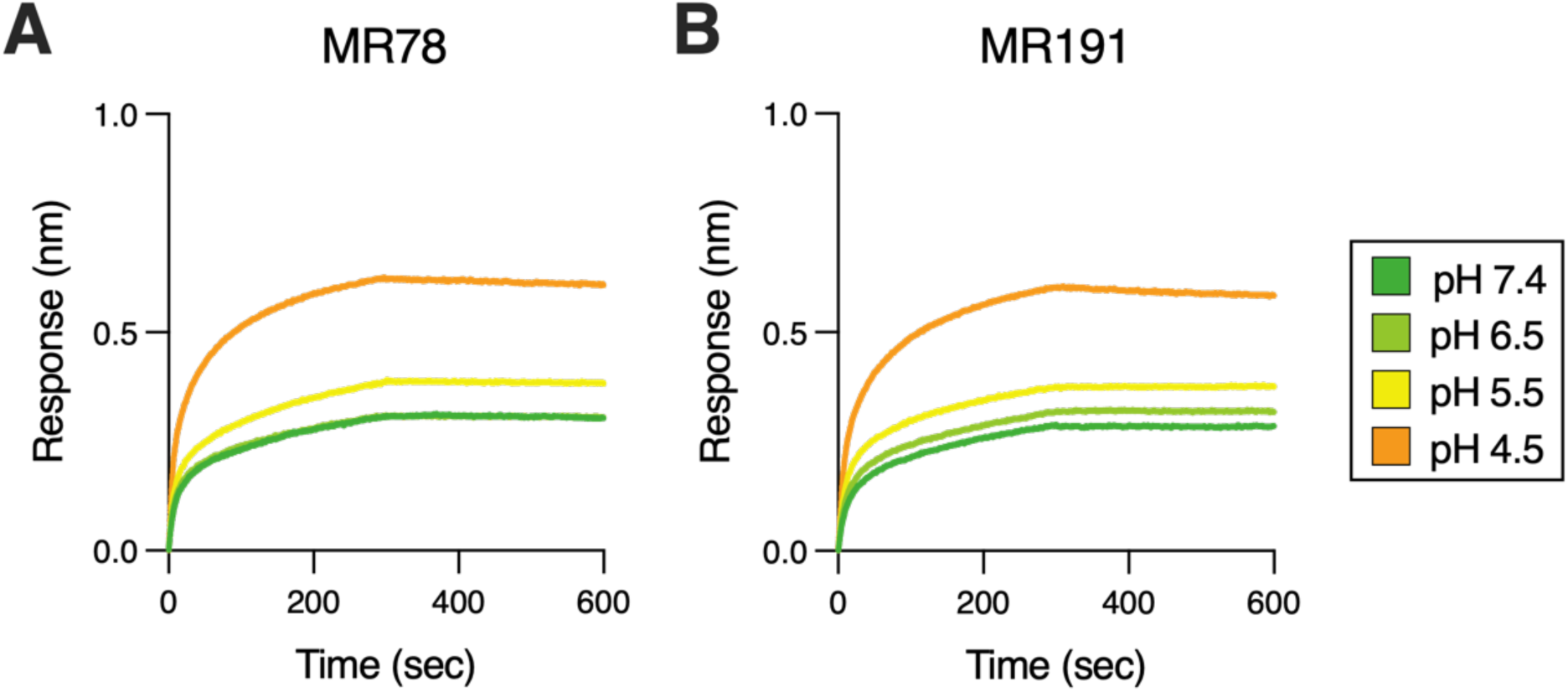
Binding of MR78 (A) and MR191 (B) IgGs at a concentration of 100 nM to immobilized MARV GPΔMuc at the indicated pH as measured by biolayer interferometry. Data are representative of two biological replicates using distinct batches of protein and antibodies.

**Figure S4.**
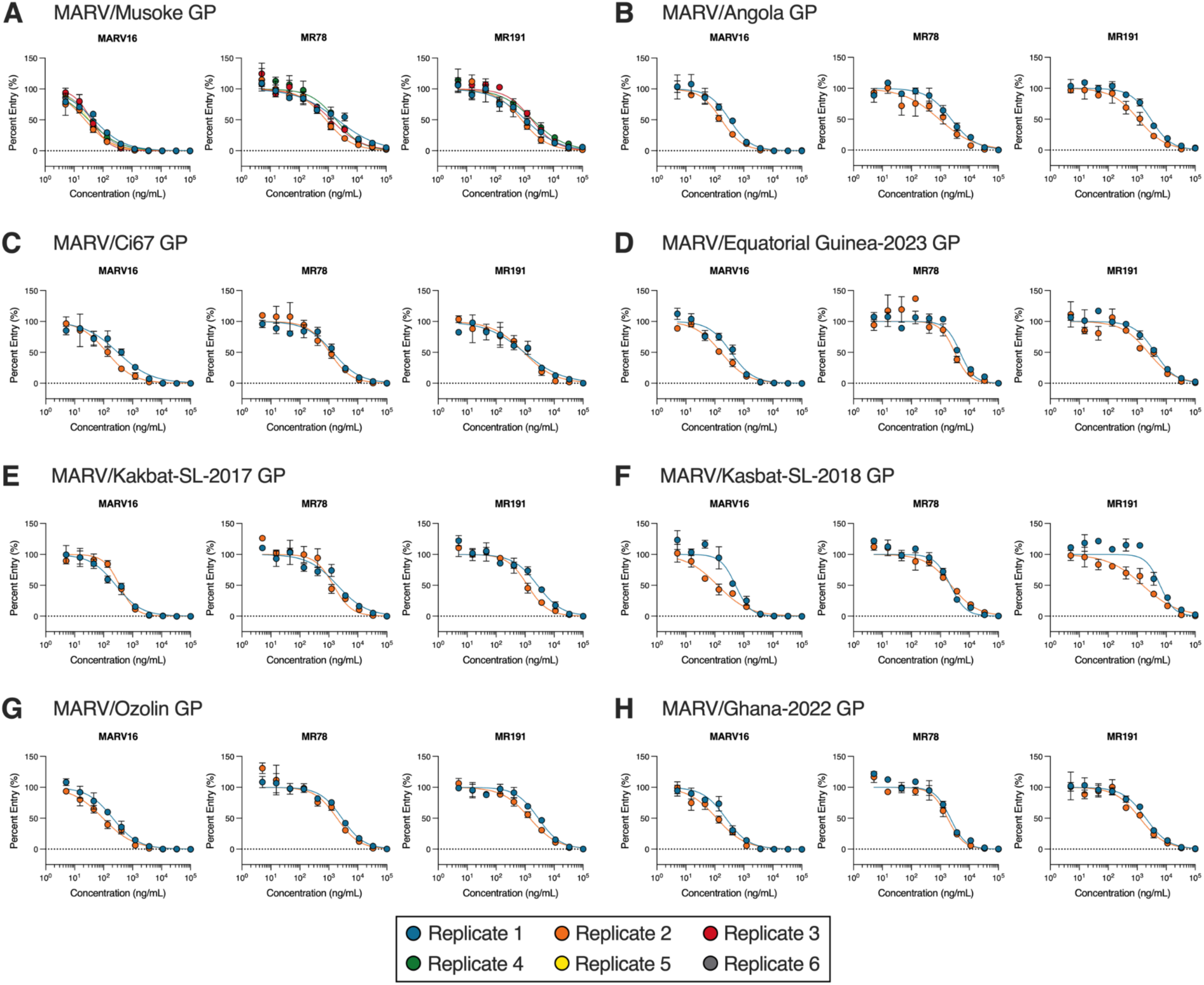
Neutralization dose-response curves for MARV16, MR78, and MR191 against VSV pseudotyped with the MARV/Musoke (A), MARV/Angola (B), MARV/Ci67 (C), MARV/Equatorial Guinea-2023 (D), MARV/Kakbat-SL-2017 (E), MARV/Kasbat-SL-2018 (F), MARV/Ozolin (G), or MARV/Ghana-2022 GP (H). Each of the two to six biological replicates used distinct batches of pseudoviruses and antibodies and data are shown as the mean ± standard error of technical triplicates.

**Figure S5.**
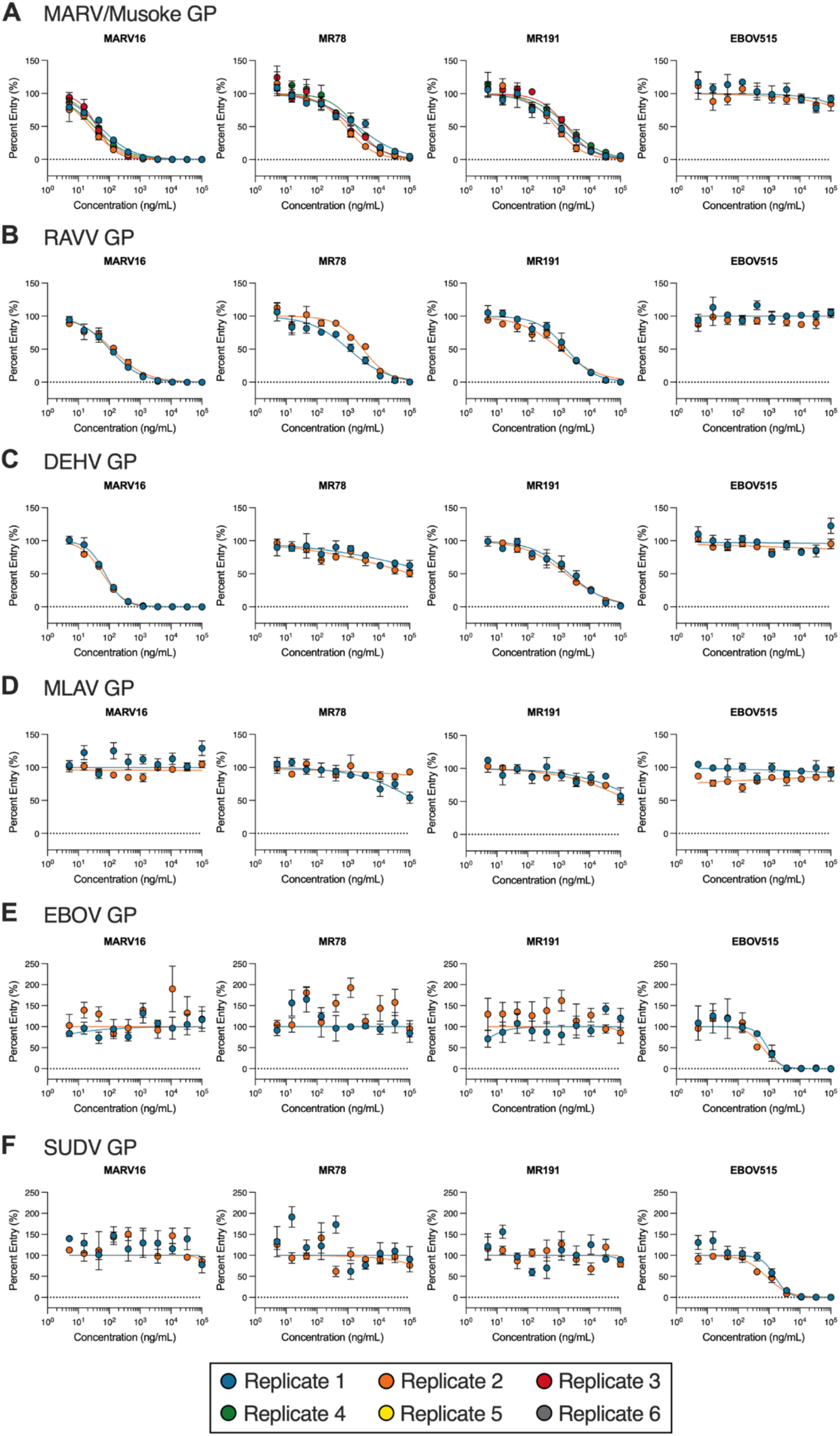
Neutralization dose-response curves for MARV16, MR78, MR191, and EBOV515 against VSV pseudotyped with the MARV/Musoke (A), RAVV (B), DEHV (C), MLAV (D), EBOV (E), or SUDV GP (F). Each of the two to six biological replicates used distinct batches of pseudoviruses and antibodies and data are shown as the mean ± standard error of technical triplicates.

**Figure S6.**
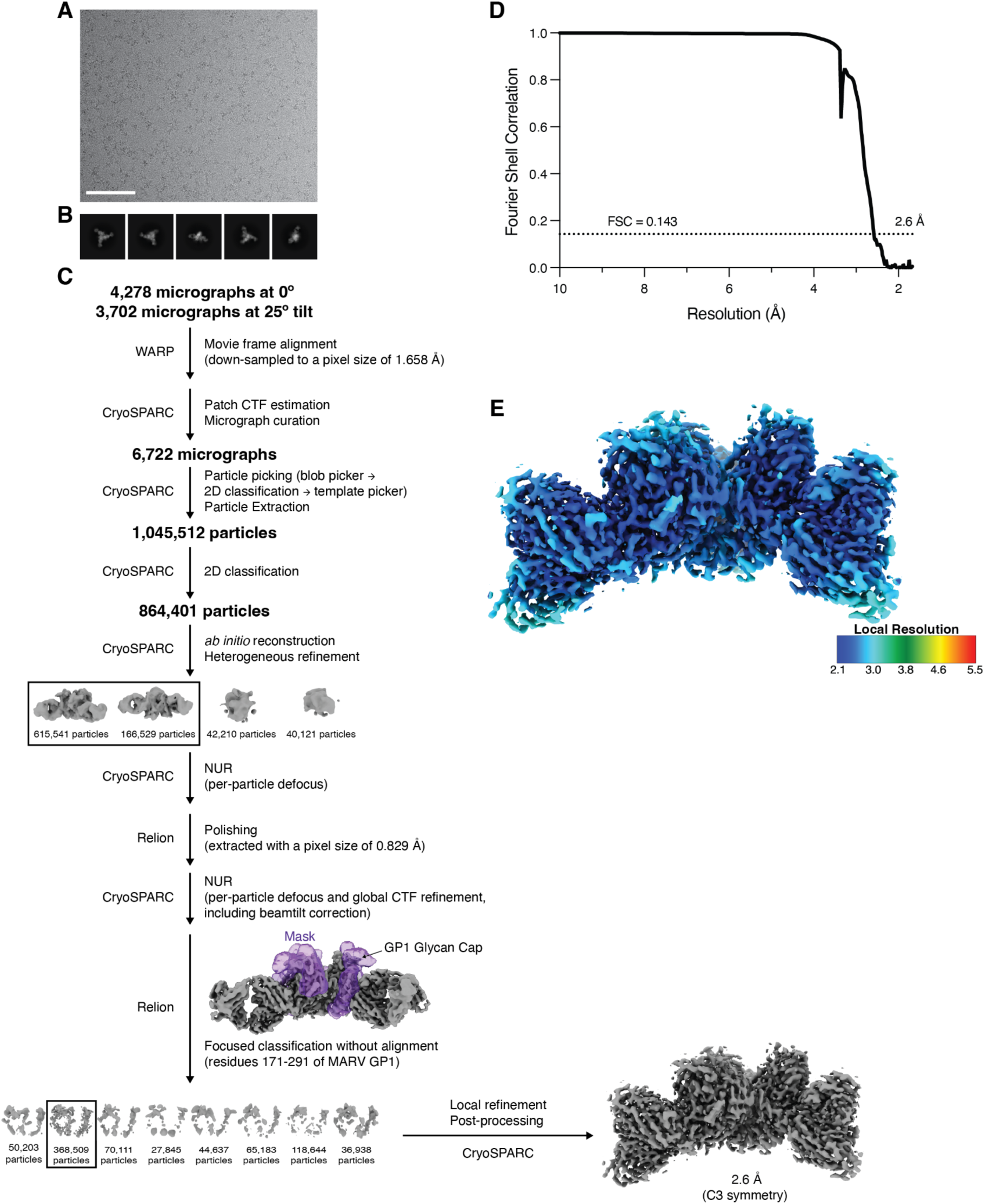
**A-B)** Representative cryo-EM micrograph (A) and 2D class averages (B) obtained for MARV GPΔMuc in complex with MARV16 Fabs. Scale bar: 100 nm. **C)** Cryo-EM data processing workflow. **D)** Gold-standard fourier shell correlation curves for the MARV GPΔMuc-MARV16 complex. **E)** Local resolution calculated with cryoSPARC for the locally refined MARV GPΔMuc-MARV16 complex (local filter map).

**Figure S7.**
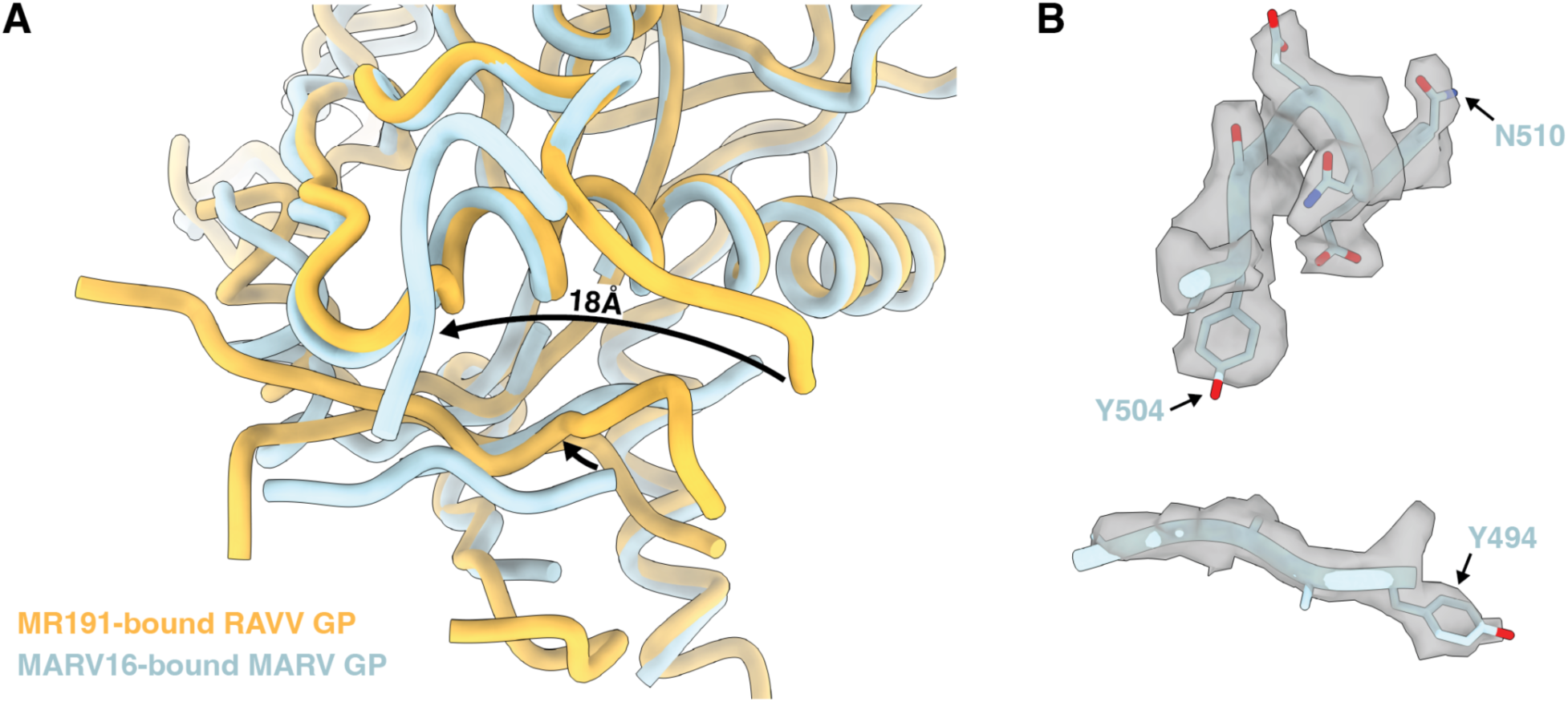
**A)** Superimposition of the MR191-bound RAVV GP (PBD: 6BP2; orange) and MARV16-bound MARV GP (blue) comparing the GP2 wing domain of the two models. The arrows indicate the position of the wing residues in each model. The MR191 and MARV16 Fabs are hidden for clarity. **B)** The MARV GP2 wing modeled into the cryo-EM density map (gray surface).

**Figure S8.**
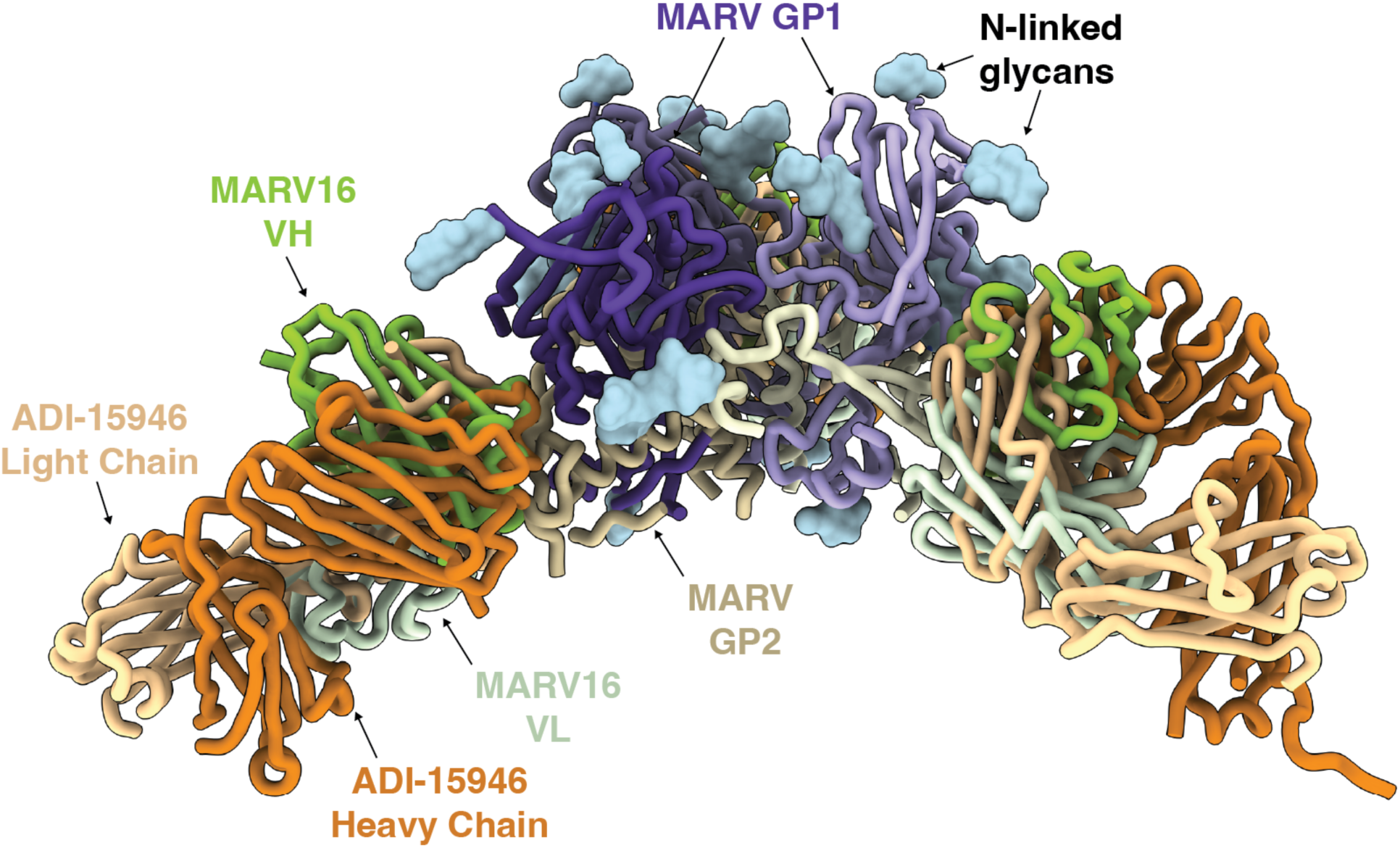
Comparison of the binding modes of MARV16 (green) and the pan-ebolavirus antibody, ADI-15946 (orange). The EBOV GP trimer from the EBOV GP-ADI-15946 complex structure(PDB: 6MAM) was superimposed with the MARV GP trimer from the MARV GPΔMuc-MARV16 structure to compare the ADI-15946 and MARV16 binding poses. MARV GP1 and GP2 are shown in different shades of purple and beige, respectively. N-linked glycans are rendered as cyan surfaces.

**Figure S9.**
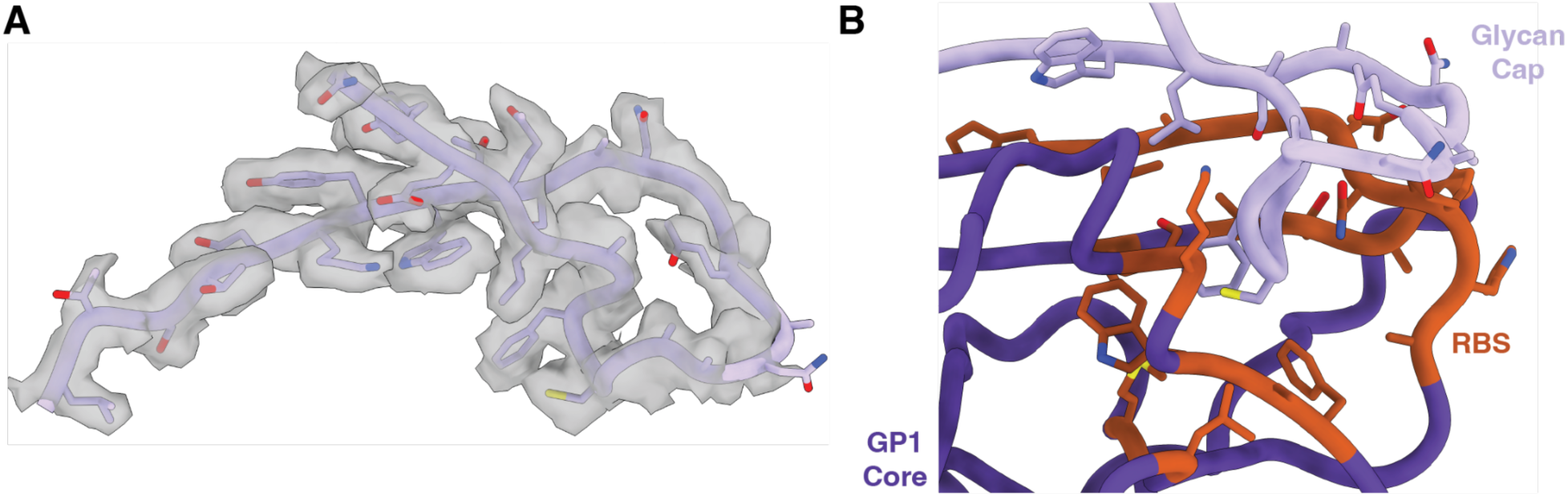
**A)** The MARV GP1 glycan cap (residues 191-219) modeled into the cryo-EM density map (gray surface). **B)** View of the putative RBS residues (shown in orange) that are shielded by the MARV GP1 glycan cap (light purple).

**Table S1.**
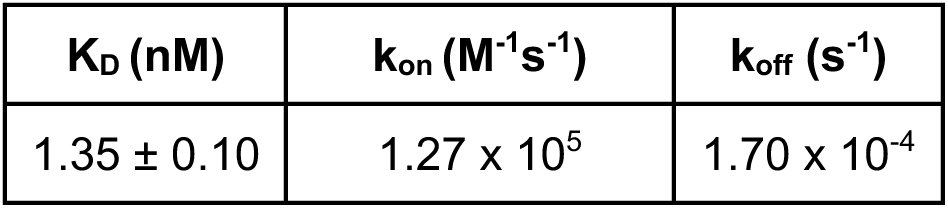
Binding kinetics of the MARV16 Fab to immobilized MARV GPΔMuc. Values are presented as mean ± standard deviation obtained from two biological replicates using distinct batches of protein.

| **K<sub>D</sub> (nM)** | **k<sub>on</sub> (M<sup>-1</sup>s<sup>-1</sup>)** | **k<sub>off</sub> (s<sup>-1</sup>)** |
| --- | --- | --- |
| 1.35 ± 0.10 | 1.27 × 10<sup>5</sup> | 1.70 × 10<sup>-4</sup> |

**Table S2.**
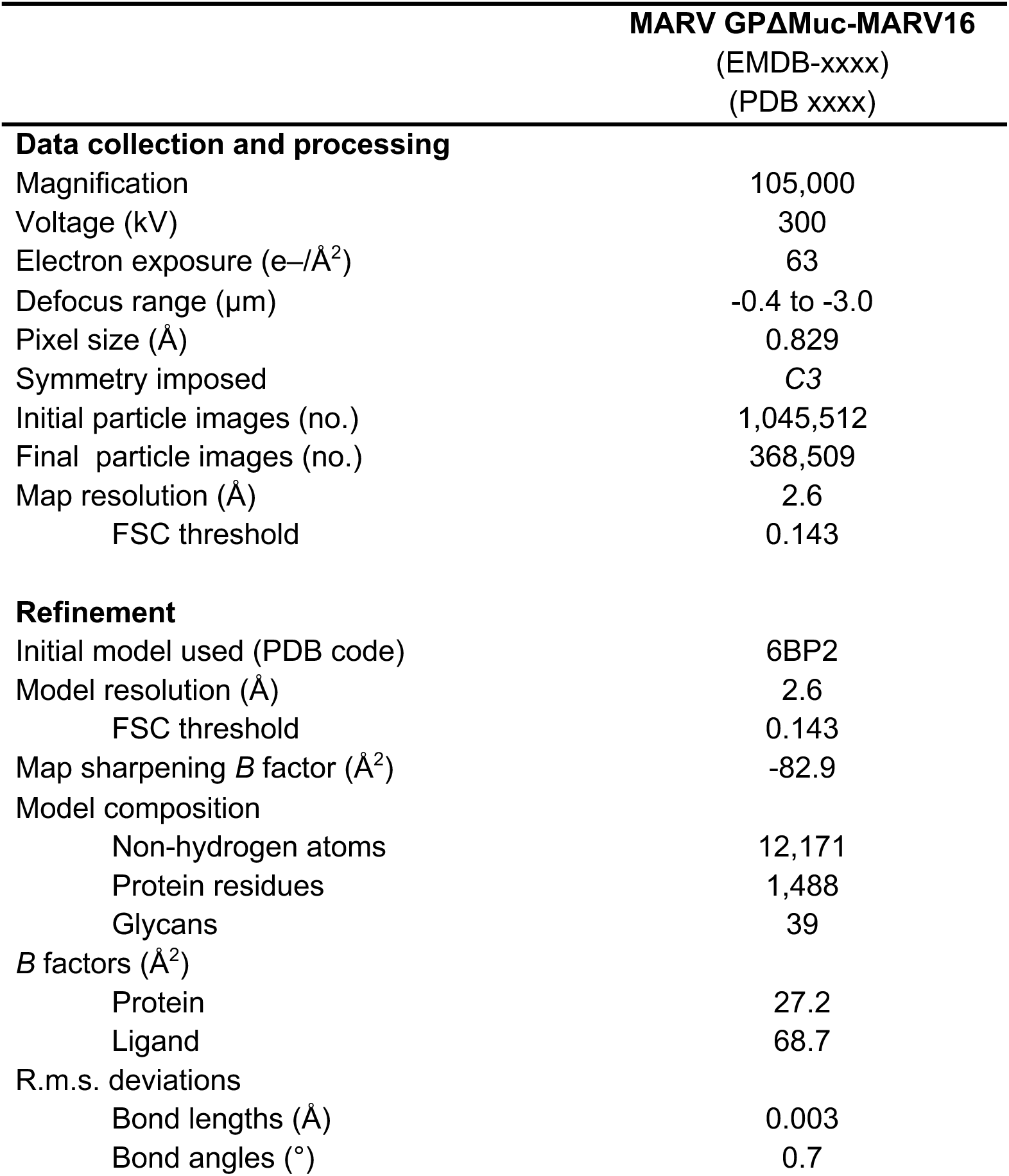
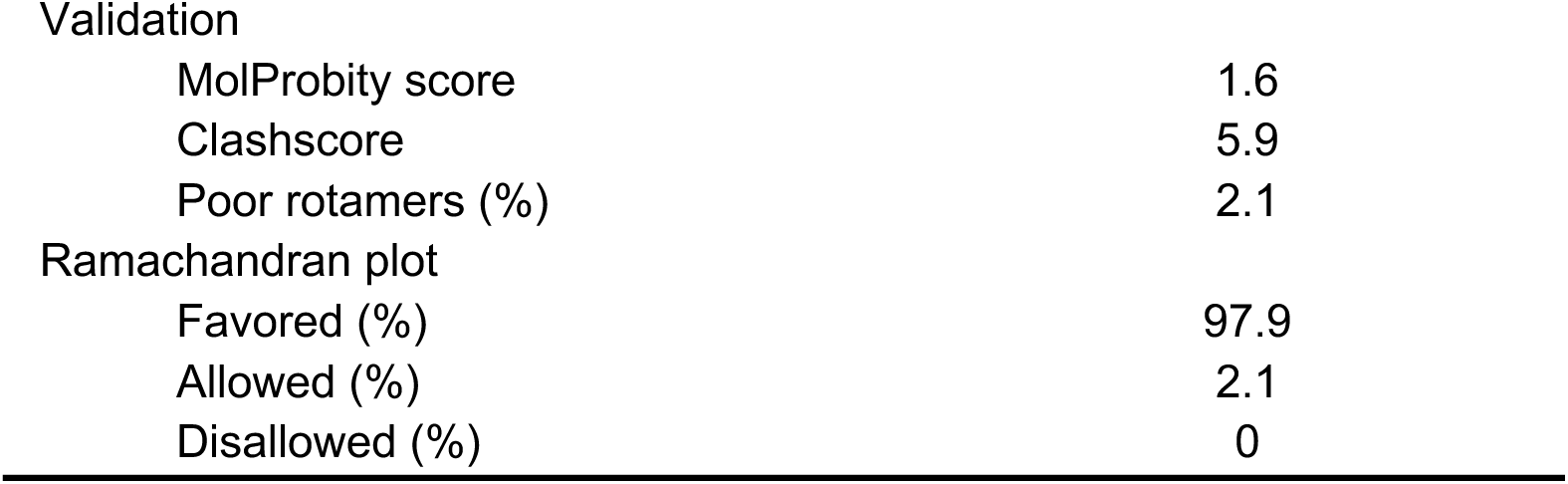
Cryo-EM data collection, refinement and validation statistics.

